# Sequenced-based GWAS for linear classification traits in Belgian Blue beef cattle reveals new coding variants in genes regulating body size in mammals

**DOI:** 10.1101/2023.06.27.546701

**Authors:** JL Gualdron Duarte, C Yuan, AS Gori, GCM Moreira, H Takeda, W Coppieters, C Charlier, M Georges, T Druet

**Affiliations:** Unit of Animal Genomics, GIGA-R & Faculty of Veterinary Medicine, University of Liège, Avenue de l’Hôpital, 1, 4000 Liège, Belgium; Walloon Breeders Association, Rue des Champs Elysées, 4, 5590-Ciney, Belgium; GIGA Genomic platform, GIGA-R, University of Liège, Avenue de l’Hôpital, 1, 4000 Liège, Belgium

## Abstract

Cohorts of individuals that have been genotyped and phenotyped for genomic selection programs offer the opportunity to better understand genetic variation associated with complex traits. Here, we perform an association study for traits related to body size and muscular development in intensively selected beef cattle. We leveraged multiple trait information to refine and interpret the significant associations. After a multiple-step genotype imputation to the sequence-level for 14,762 Belgian Blue beef (BBB) cattle cows, we performed a GWAS for 11 traits related to muscular development and body size. The 37 identified genome-wide significant QTL could be condensed in 11 unique QTL regions based on their position. There was evidence for pleiotropic effects in most of these regions (e.g., correlated association signals, overlap between credible sets of candidate variants – CSCV). We consequently applied a multiple-trait approach to combine information from different traits to refine the CSCV. In several QTL regions, we identified strong candidate genes known to be related to growth and height in other species such as *LCORL-NCAPG* or *CCND2*. For some of these genes, relevant candidate variants were identified in the CSCV, including three new missense variants in *EZH2*, *PAPPA2* and *ADAM12*, possibly two additional coding variants in *LCORL*, and candidate regulatory variants linked to *CCND2* and *ARMC12*. Strikingly, four other QTL regions were related to five (recessive) deleterious coding variants previously identified. Heterozygotes for several of these mutations have favorable effects for muscular development traits. Our study further supports that a set of common genes controls body size across mammalian species. In particular, we added new genes to the list of those associated with height in both human and cattle. We also identified new strong candidate causing variants in some of those genes, strengthening the evidence of the causality of these genes. Several breed-specific recessive deleterious variants were identified in our QTL regions, probably as a result of the extreme selection for muscular development in BBB cattle.

## Background

Reference populations built to implement genomic selection [1] in livestock species are valuable resources to understand the genetic basis for variation in complex traits. The size of these cohorts of genotyped and phenotyped individuals are increasing through the years, while genomic selection is applied to more and more livestock species and breeds [e.g., 2, 3]. Although data collection focuses mainly on traits of agronomic importance, recorded phenotypes might also help to study complex traits of interest for other applications. For instance, these populations can be used to study traits related to health such as fertility (in particular, in the context of artificial reproductive technologies that are massively used in livestock), or to fundamental biological processes such as meiotic recombination [e.g., 4, 5]. Traits recorded in multiple species allow to understand to which extent the genetic architecture of complex traits is conserved across species or how it evolves [e.g., 6]. A typical example would be the study of stature in mammals, where results obtained in human, dog, cattle and horse have already been compared, revealing that genome-wide association signals are enriched in genes associated in other species [e.g., 6–8]. Indeed, association studies and scan for signatures of selection identified genes associated with height in multiple species as *IGF1*, *PLAG1* or *LCORL-NACPG* [e.g., 7, 9–13]. Similarly, variations in *MSTN* affecting muscular development have been identified in several species including cattle, sheep, dog, horses, pig and human [11, 12, 14–16]. Interestingly, livestock provides information on these complex traits in populations under intensive selection and with reduced effective population size.

The Belgian Blue beef (BBB) cattle breed represents an example of a breed intensively selected for muscular development. As a result, a loss-of-function mutation in the myostatin gene causing the double muscling phenotype [14] has been fixed through selection [14, 17]. However, additional genetic variation for muscular development is still available within the breed and has been exploited to further increase this trait [see 17]. In addition, several recessive deleterious variants under balancing selection were segregating in the population at high frequency before the implementation of genetic tests, including a 2-bp deletion in the open reading frame of *MRC2* [18] and a splice-site variant in *RNF11* [19]. For these loss-of-function (LoF) variants, heterozygote advantage resulted from the favorable effect of these alleles on selected traits such as muscular development or stature. Similarly, a R844Q missense variant in *WWP1* presented evidence for both a recessive deleterious effect and a favorable effect on muscular development [20]. Indeed, significantly fewer homozygotes than expected were observed in the population despite a relatively high frequency, indicating selection against homozygotes and a recessive effect. In the last years, a genomic selection program has been implemented in BBB cattle, the genotyping of individuals starting in 2016. The reference population is phenotyped mainly for a set of linear classification traits, related to body size and muscular development, that are routinely recorded on adult females. This population represents an example of a cattle breed intensively selected for muscular development and for body size to a lesser extent.

Genome-wide association studies (GWAS) are one of the main tools to decipher the architecture of complex traits. Reference populations offer the opportunity to apply such scans in livestock species, but the marker density is often too low to capture well all causative variants and to perform the fine-mapping. Therefore, genotype imputation to the sequence level thanks to a reference population [21, 22] is a recommended practice. Such sequence-based GWAS have already been successfully implemented in cattle [e.g., 8, 23–25]. In addition to sequence-based level approaches, multiple traits association methods [26, 27] might be useful to improve the fine-mapping resolution since subsets of recorded traits are often genetically correlated and affected by shared pleiotropic variants.

We herein performed such a sequence-based GWAS in BBB cattle for linear classification traits related to body size and muscular development. Multiple-traits information was leveraged to refine the set of candidate variants and to interpret the relation between associations for different quantitative trait loci (QTL) mapping to the same genomic region. For several of the identified QTL, we found genes associated to stature in other species among the candidate genes, providing further support for the presence of common genes regulating body size in mammals. For several of them, we identified coding variants as strong candidates that give stronger evidence of the causality of these genes. Besides these QTL, other were associated to five recessive deleterious variants having favorable effects at heterozygous state on traits related to muscular development.

## Material and methods

### Data

Several groups of BBB cattle were used in the present study. The first group consisted in a set of 14,762 cows having both genotypes and phenotypes, and will be referred to as the “mapping population”. These cows were genotyped on six different versions of the Illumina Bovine Low Density genotyping arrays used by the EuroGenomics consortium (from 9983 to 20,502 SNPs), on EuroGenomics MD arrays (three versions with 51,809 to 57,979 SNPs), or on the Illumina Bovine50K medium density arrays (two versions, with 54,001 and 54,609 SNPs). Linear classification scores (assessed visually by a technician) for 10 traits including length, pelvis length, height, chest width, pelvis width, rib shape, rump, top muscling, shoulder muscling and buttock muscling (side and rear view) were available for 14,476 cows. In addition, height was measured at withers for 12,904 cows. These phenotypes were corrected for fixed effects from the genetic evaluation to obtain trait-deviations (TD). In addition, a set of 717 AI bulls was genotyped on the BovineHD genotyping array. Among these, 658 were also genotyped on the LD. Another group of 199 animals, including 66 AI bulls, was genotyped on both Illumina BovineLD and Bovine50K. A fourth group of 9502 individuals without phenotype were genotyped on EuroGenomics MD arrays. These groups of individuals were available as reference populations for genotype imputation. The number of individuals genotyped on the different arrays is described in Additional File 1: Table S1. Finally, whole-genome sequence data was available for 230 bulls at an average coverage depth of 35x, ranging from 11x to 68x.

### Read mapping and variant calling procedure

The sequencing data came from two distinct experiments. For a first group of 50 bulls previously described in Charlier et al. [20], DNA was extracted from sperm using standard procedures. PCR-free libraries were sequenced at the CNAG in Barcelona on an Illumina HiSeq 2000 with a paired-end protocol (2 x 100 bp), each sample being sequenced on multiple lanes. For the 180 remaining bulls, DNA was extracted from blood or sperm and paired-end sequencing (2 x 150 bp) was performed on an Illumina NovaSeq 6000 sequencer. Reads were aligned to the ARS-UCD1.2 (BosTau9) bovine genome assembly [28] with BWA-MEM v0.7.5a [29], sorted with Sambamba v0.6.6 [30] and PCR duplicates were marked with Picard Tools v2.7.1 (https://broadinstitute.github.io/picard/). BAM files were recalibrated using GATK4 v4.1.7.0 [31], using a list of 110,270,194 known variants provided by the 1000 Bull Genomes project [32] as known polymorphic sites. Recalibrated BAM files from samples sequenced on different lanes at the CNAG were then merged per bull. Individual variant calling was performed with HaplotypeCaller (GATK4 v4.1.7.0) and the joint genotyping of all the Genomic Variant Call Format (GVCF) files was subsequently done with GenotypeGVCFs (GATK4 v4.1.7.0) in 5 Mb windows. Quality scores from the resulting VCF were then recalibrated using VariantRecalibrator (GATK4 v4.1.7.0). A set of 1,213,314 SNPs from commercial genotyping arrays [33] was used as truth set, and the ∼110 M SNPs provided by the 1000 Bull Genome (see above) project as known set. This procedure defines quality thresholds that would result in the conservation of different fractions of the truth set (e.g., 90, 95, 97.5%). Variants with a quality score below the 97.5 threshold, with a MAF < 0.01, and multi-allelic sites were filtered out, resulting in a set of 12,830,339 SNPs and 2,502,613 indels.

### Marker selection and genotype imputation

Genotype imputation from low-density to the sequence level was performed in successive steps [34], using medium density (MD) and high-density (HD) levels as intermediate steps. The LD level consisted in all cows from the mapping populations genotyped on LD arrays (11,521 cows). The reference population at the MD level consisted in i) cows from the mapping population and other individuals genotyped on EuroGenomics MD arrays (12,475 animals) or genotyped on both LD and Bovine50K arrays (467 individuals) and ii) AI bulls genotyped simultaneously on LD and bovineHD arrays (658 bulls). At the HD level, the sequenced bulls or those genotyped on the BovineHD arrays defined the reference population, corresponding to 890 unique individuals. At each level, we selected markers that were common to all involved panels (for individuals genotyped on two arrays, selected markers had to be present on at least one of them) and that were useful for the imputation procedure (shared either with the previous or with the next level). We filtered out markers with a call rate below 0.95, with a minor allele frequency (MAF) below 0.01, deviating from Hardy-Weinberg proportions (p < 0.001) or with more than 10 Mendelian inconsistencies (e.g., opposite homozygous genotypes in parent/offspring pairs), and individuals with less than 90% genotyping rate. As a result, we selected respectively 7,525, 32,318 and 611,322 SNPs at the LD, MD and HD levels. Beagle 4.1 [35] was first applied to the WGS and HD reference panels to improve genotype calls and impute sporadically missing genotypes. Target and reference panels were phased with ShapeIT4.2 [36] and Minimac4 [37] was applied to achieve the imputation in the target panel. After each intermediary imputation step, we discarded SNPs with a MAF < 0.02 in the imputed set, and those with a reported evaluation accuracy below 0.80. In addition, we removed SNPs not useful for the next imputation step (for instance, SNP shared between LD and MD arrays but absent from the HD array; these are useful for the first imputation step but no longer in the second step). As a result, we conserved 28,893 and 572,667 SNPs at the MD and HD level for imputation to the next level. These additional cleaning steps were applied to keep only SNPs that were expected to be accurately imputed. After the last imputation step, the final VCF file contained 11,537,240 SNPs and indels with a MAF > 0.01.

### Genome-wide association study

Single-trait GWAS (ST-GWAS) were performed on each trait using the following linear mixed model (LMM) approach with GEMMA [38] to test the association with marker *i*:

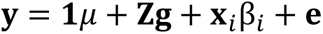

where **y** is the vector of trait deviations, µ is the overall mean effect, **g** a vector containing the random additive polygenic effects of the corresponding cows, β_i_ is the additive effect of the tested SNP, **x***_i_* is a vector containing the allele dosage for the corresponding cows at marker *i*, **e** is a vector of random error terms and **Z** is an incidence matrix indicating which animal is associated with the phenotype. The covariance structure among random polygenic effects **g** is a function of the genomic relationship matrix **G** obtained from the 28,893 SNPs from the MD level and constructed using the first method proposed by VanRaden [39]. The number of independent tests per genome-scan was estimated with the procedure described in Druet *et al.* [17]. Briefly, we performed a genome-scan for association with height using a simple regression, providing us a distribution of uncorrected p-values. We repeated then genome-scans on 100,000 random permutations of the phenotypes and recorded the best p-value for each scan, providing the distribution of best p-values under H0 that allowed us to obtain corrected p-values for the first scan. Finally, the number of independent tests was estimated to be equal to 500,900 (rounded to 500,000) based on the comparison of the uncorrected and corrected p-values and using the Sidak formula. As we repeated the association study for 11 traits, we also estimated the number of independent traits using the meff function (method = Galwey) from the poolR R package [40] and obtained a value of seven. As a result, we set the significance threshold to 1.43e-8 (-log_10_P > 7.84) after applying a Bonferroni correction for 3,500,000 independent tests. In regions where a significant QTL was detected, we also considered that other traits with significance levels below 1e-7 (-log_10_P > 7) presented evidence for association with a QTL in the region.

The set of candidate causative variants, referred to as credible sets (CS), were defined with two approaches. First, an Iterative Bayesian Step-wise Selection (IBSS) approach implemented in SuSiE [41, 42], relying on summary statistics obtained from the ST-GWAS and on the LD pattern among SNPs, was applied to identify a CS of SNPs with a probability > 0.95 of containing the causal variable (based on the individual Posterior Inclusion Probabilities (PIP) from each SNP). Note that this approach also provides multiple CS when several independent effects are detected in the QTL region. In addition, LD-based CS were obtained by selecting all the SNP in high LD with the lead variants (with the minimal LD level r² set to 0.90 or 0.80).

### Comparison of associations across traits and multiple-trait association studies

For each trait, we identified genome-wide significant associations and considered other significant associations within 1-Mb as part of the same QTL region (QTLR). When significant SNPs were identified for different traits less than 1-Mb apart, the QTLR were considered as a single QTLR. To investigate whether associations for different traits in the same QTLR resulted from a pleiotropic QTL or from closely linked QTL, we compared the SNP significance levels obtained for pairs of traits. To that end, we selected all the SNPs 1 Mb around the lead SNP and computed correlations among t-values or signed p-values (on a –log10 scale) to take into account the effect direction [e.g., 43, 44]. Furthermore, we excluded non-significant signals (p > 0.05) as effects are not expected to be shared for these unassociated SNPs [44]. In addition, we studied overlap between CS obtained for all the traits presenting significant association in the QTLR to find further evidence of pleiotropy.

As we found evidence that several associations were shared across multiple traits, we decided to run multiple-trait GWAS (MT-GWAS) with the Multivariate LMM approach implemented in GEMMA [45]. As recommended, we limited the number of phenotypes in the multivariate analyses. Therefore, we run the model on two group of six traits selected based on shared associations. The first group contained traits related mainly to body size (height, length, pelvis length, pelvis width, chest width, rib shape), whereas the second was more related to muscular development (shoulder and top muscling, side and rear-view of buttock muscling, rump and chest width). This MT-GWAS was used to combine information from multiple traits to improve the fine-mapping by defining multiple-trait LD-based CS. The MT-GWAS information was considered for fine-mapping when at least one of the six traits presented evidence of association (-log_10_P > 7). These correspond to association levels that would be genome-wide significant in a single-trait GWAS. When both MT-GWAS matched this condition, their CS were merged.

### Annotation of associated variants

In each CS, we search for candidate causal variants. In addition to the statistical evidence, we relied on the annotation of the variants in the CS obtained with Variant Effect Predictor (VEP) v95.0 [46] that provides also the predicted impact (from MODIFIER to HIGH) and the SIFT score for missense variants. PhastCons conservation scores across 30 vertebrates [47] and GERP scores across 91 mammals [48] were used as conservation metrics. Furthermore, information available from the literature was considered for variants previously reported. Finally, we investigated whether SNP in the CS overlapped with core and consensus segments called from ATAC-seq peaks in a recently released catalogue (Yuan et al., 2023), available at http://tinyurl.com/2p8kr7uv, or with credible sets of blood and liver cis-eQTL reported in the same study.

### Conditional mapping

Subsequently, one candidate variant was selected per CS to perform a conditional mapping scan by fitting them as fixed effects in the LMM. For each QTLR, this conditional mapping allows to determine how well the tested candidate causal variant captures the QTL signal and whether it captures the signal for different traits, providing eventually further evidence for causality and pleiotropy. In addition, it allows to determine whether a single or multiple QTL affect the same trait in the QTLR. The conditional mapping was performed in eleven 10 Mb regions centered around the lead variants and encompassing a total of 516,465 SNPs (Additional File 1: Table S2). Using the same approach as before, we estimated that the number of independent tests was approximately equal to 6000. For each QTLR, the conditional analyses were performed only for traits presenting evidence for association in the first GWAS (-log_10_P > 7), resulting in approximately 3.1 independent traits per region on average. As a result, we set the significant threshold to 2.5e-6 (-log_10_P > 5.6) to account for a total of approximately 20,000 independent tests. We repeated the same procedure if new significant QTL were detected in the QTLR.

## Results

### Identification of eleven QTL regions affecting multiple traits

Application of the ST-GWAS for eleven distinct traits resulted in the identification of 37 QTL (Figure 1, Additional File 1: Table S3). The most significant QTL (p < 1e-15) were located on Bos taurus autosomes (BTA) 5, 6 and 14 and mainly associated with traits such as height or body length (see Figure 2). Based on their position, the QTL could be organized in 11 groups (less than 1Mb distance between peaks). Most of the QTLR affected multiple traits (Figure 1), with QTLR on BTA14 and BTA19 harboring significant associations with respectively 8 and 7 traits. In these QTLR, association levels or t-values obtained for different traits were often highly correlated (Additional File 2). This is illustrated in Figure 3 for the QTLR on BTA14 (see Additional File 3: Figures S1-S8 for other traits). In general, effects for traits such as height or length were negatively correlated with effects estimated for muscular development traits. In addition, the credible sets (CS) of candidate variants identified for different traits were overlapping for several QTLR. For instance, the SuSiE CS from all traits with a significant QTL shared at least one variant for eight QTLR (out of nine QTLR associated to two or more traits – Additional File 1 – Table S4). For three QTLR, the CS were even identical across all traits. Similar results were obtained when using LD-based CS (LDCS) obtained by selecting all SNP with a r² > 0.80 with the lead variants (slightly fewer sharing with a threshold of r² > 0.90). Both approaches pointed to similar CS. For instance, LDCS (r² > 0.80) and SuSiE CS obtained shared at least one common candidate variants for 34 out of 37 QTL, and the SuSiE CS was totally included in the LDCS for 29 QTL (Additional File 1: Table S5). Overall, there is thus strong evidence that several of the QTLR harbor pleiotropic variants and we decided consequently to combine multiple-trait information to define MT LD-based CS. This MT mapping was performed in two groups of six traits related either to body size or to muscular development (see list in material and methods). The resulting MT LD-based CS are described in Table 1. The median number of SNP included in these MT LD-based CS was equal to 11 (ranging from 1 to 116; mean = 30), and their span ranged from 1 bp to 1.7 Mb (median = 268 kb). Only one or two associated genes (i.e., genes with coding, intergenic or up/down stream variants in the CS) were generally found in these CS (only two regions with > 3 associated genes).

**Figure 1.**
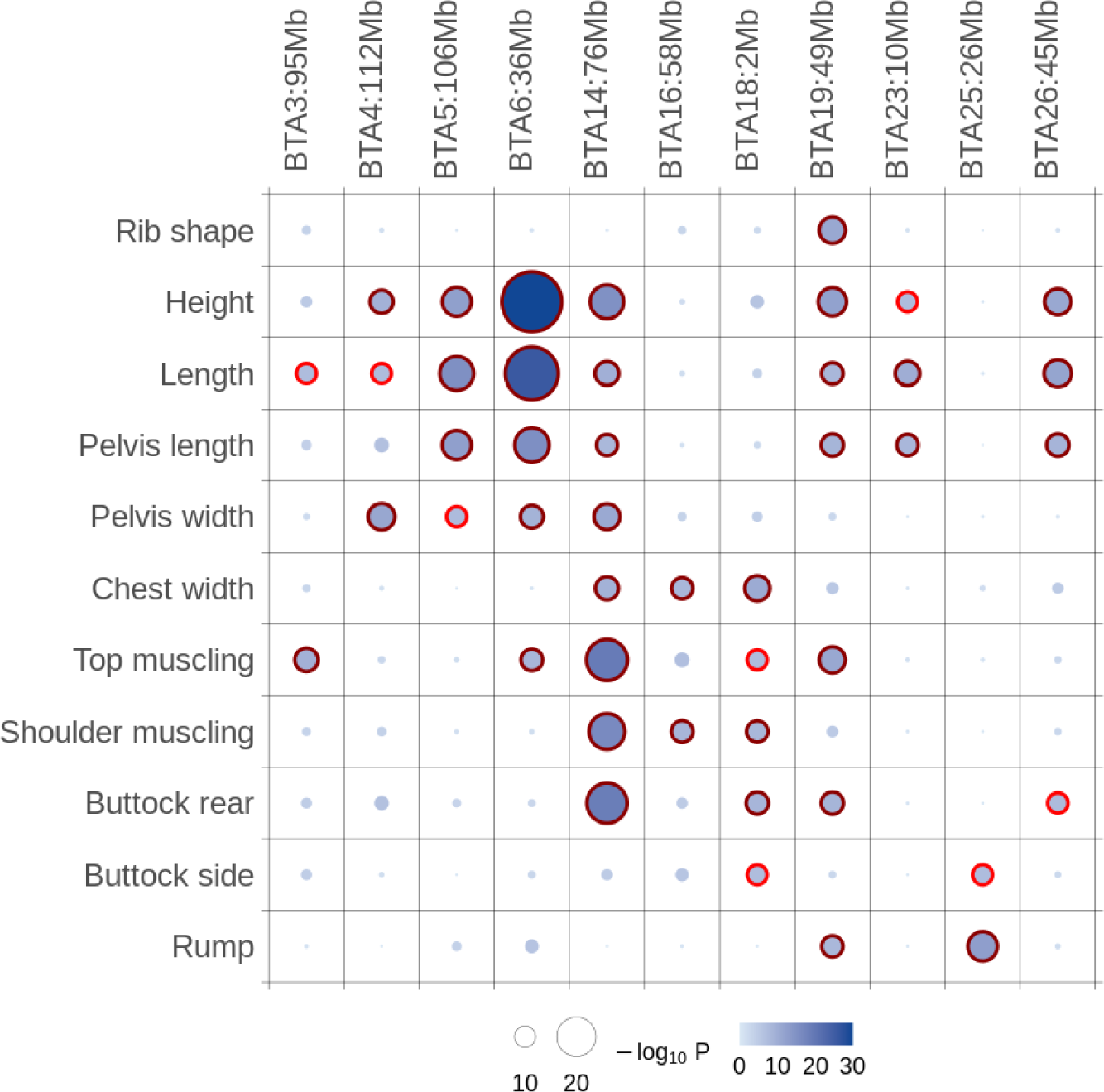
Summary of identified QTL regions. QTLR are labelled according to the corresponding chromosome and the position (rounded in Mb). The maximum significance level for each trait per QTLR is indicated by the size and color intensity of the circles. Significant QTL are indicated with dark red outer circles, whereas red outer circles are used for QTL reaching significance levels for a single genome scan.

**Figure 2.**
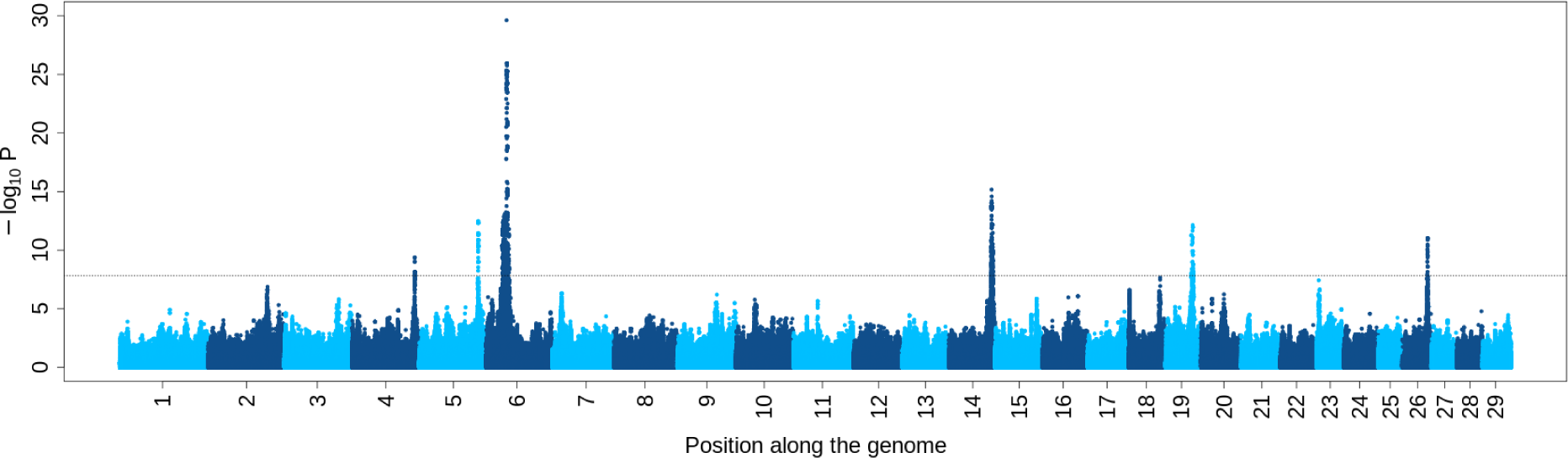
Manhattan plot for association with height. The horizontal line represents the significance level after correction for multiple testing.

**Figure 3.**
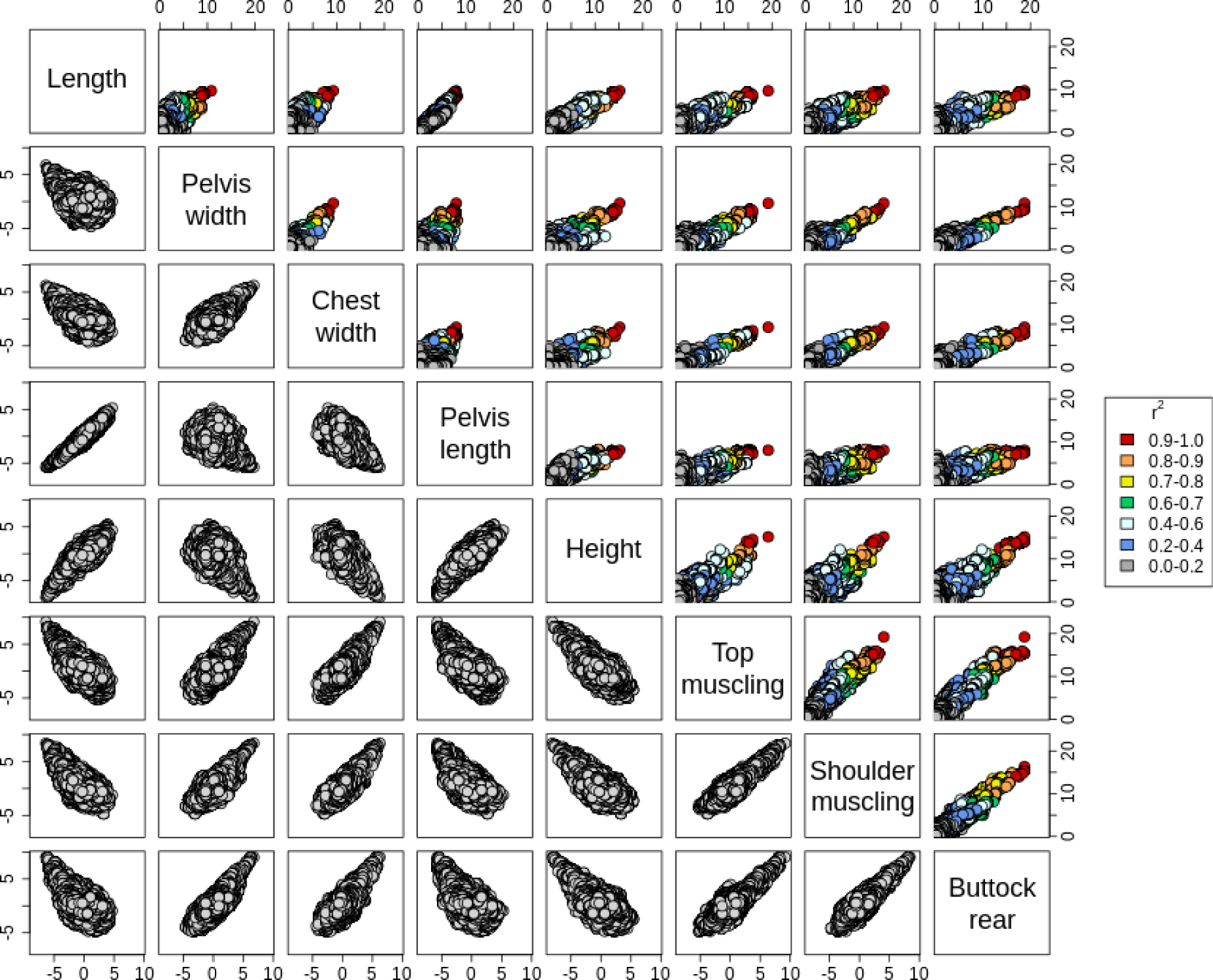
Scatterplots with association levels for different traits for the QTL region on BTA14. The selected traits are those harboring a significant signal in the QTLR. Upper diagonal: scatterplots with p-values on a negative log10 scale. The color represents the LD level with the lead SNP (from the trait with the strongest association). Lower diagonal: scatterplots with signed t-values.

**Table 1.**
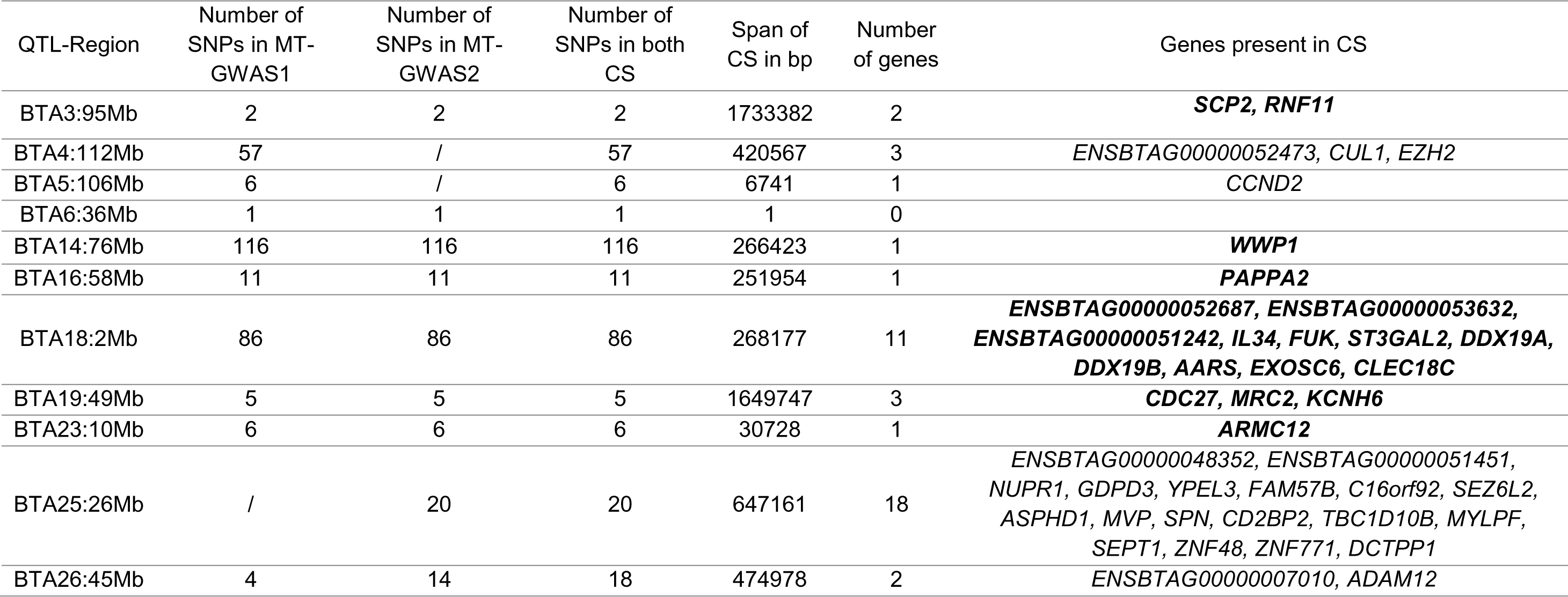
LD-based CS (r² > 0.90) identified by MT-GWAS. MT-GWAS1: MT-GWAS with traits related to height and body dimensions; MT-GWAS2: MT-GWAS with traits related to muscular development. Genes present in CS from both MT-GWAS are indicated in bold (other are specific to a single MT-GWAS).

### Four QTLR are associated to recessive deleterious coding variants

Four of the QTLR were associated to recessive deleterious variants previously identified in BBB cattle (see Table 2), three of which causing genetic defects when homozygote [18, 19, 49]. These variants were indeed present in the MT LD-based CS (Figure 4 – Additional File 3: Figure S9), and the LoF variant in *RNF11* associated to dwarfism, the 2 bp-deletion in *MRC2* associated to the crooked-tail syndrome (CTS) and the missense variant in *WWP1* reported in Charlier et al. [20] were even the lead SNP in both MT-GWAS (Additional File 1 – Table S6). The variants in *RNF11* and *WWP1* were also the lead variants in all but one of the ST GWAS. The variant in *MRC2* was the lead SNP in three out of seven ST GWAS and was included in three ST-LD based CS when the LD threshold was set to 0.90 (six if the threshold was relaxed to 0.80). Remarkably, the three variants were always present in the SuSiE CS and the variants associated with dwarfism and the CTS had always a PIP > 0.95 (*i.e.* the CS contained thus only this variant). The statistical evidence supporting these candidate variants is thus strong. Finally, the variant in *ATP2A1* associated with congenital muscular dystrophy [49] presented a LD of 0.88 with the lead SNP in both the ST GWAS for rump and the MT GWAS for muscular development traits (Additional File 3: Figure S9). To note, this variant segregates now at low frequency in the population (f = 0.011) and achieves significances only for one trait. Overall, the analysis of these previously identified variants shows that the MT LD-based approach is efficient and improves the resolution of the ST LD-based approach.

**Table 2.** Annotation of candidate or lead variants for the 11 QTLR. The frequency is reported for the alternate allele and SIFT scores are provided for missense variants.

| BTA | Position | REF | ALT | Freq | Gene | Consequences | SIFT score | GERP score | Phastcons |
| --- | --- | --- | --- | --- | --- | --- | --- | --- | --- |
| 3 | 95015373 | T | C | 0.050 | *RNF11* | Splice site variant |  | 0.862 | 0.976 |
| 4 | 112030024 | T | C | 0.470 | *EZH2* | Missense variant I549M | 0.03 | 0.222 | 1 |
| 5 | 105769735 | C | T | 0.700 | *CCND2* | Regulatory (ATAC-Seq, eQTL) |  | -4.2 | 0 |
| 6 | 36226849 | A | AT | 0.050 |  | Intergenic variant |  | 0.078 | - |
| 14 | 76227910 | C | T | 0.140 | *WWP1* | Missense variant R844Q | 0.00 | 0.530 | 1 |
| 16 | 57725284 | C | A | 0.720 | *PAPPA2* | Missense variant P282T | 0.00 | - | 1 |
| 18 | 1673649 | A | AT | 0.270 |  | Regulatory (ATAC-Seq) |  | 0.149 | - |
| 19 | 47095175 | CAG | C | 0.030 | *MRC2* | Frameshift variant |  | 0.504 | 0 |
| 23 | 9716619 | G | A | 0.480 | *ARMC12* | Regulatory (ATAC-Seq, eQTL) |  | -0.678 | - |
| 25 | 25933247 | G | A | 0.010 | *ATP2A1* | Missense variant R559C | 0.00 | 0.222 | 0.984 |
| 26 | 45553105 | G | A | 0.590 | *ADAM12* | Missense variant A582V | 0.03 | 1.220 | 0.890 |

**Figure 4.**
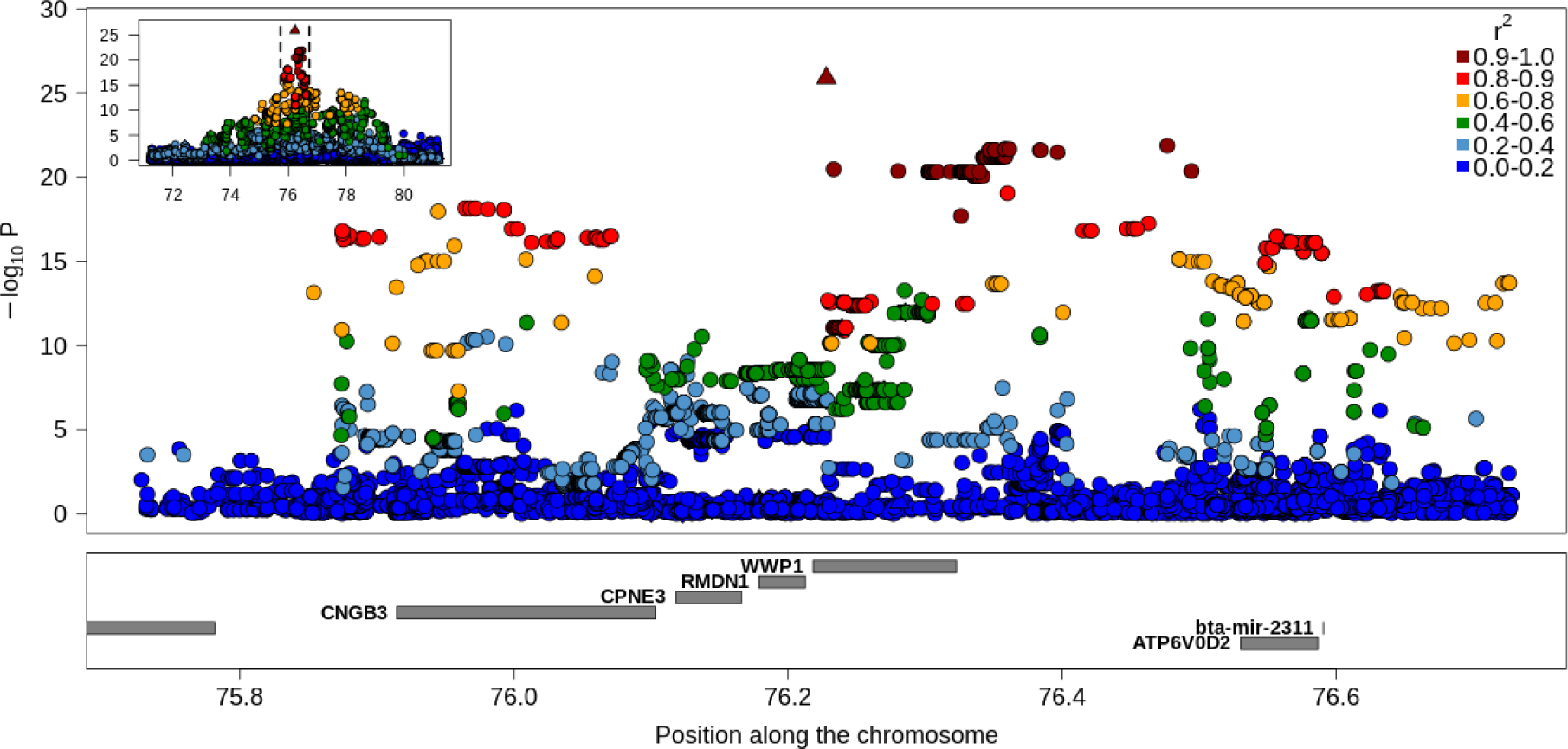
Regional association plot for the QTLR on BTA14. The results correspond to the MT-GWAS with traits related to body size. The colors represent the LD level with the lead variant and the symbols indicate the predicted impact of the variant (● modifier, ♦ low impact, ▴moderate impact, ▪ high impact). The positions of the genes are in the lower track.

### Identification of three new missense variants in genes associated with growth disorders in other species

For three additional QTLR, the MT LD-based CS contained non-synonymous variants in genes that have been previously associated to human height or growth (see the discussion for more details). These include a I549M missense variant in *EZH2* (QTLR on BTA4: Figure 5), a P282T missense variant in *PAPPA2* (QTLR on BTA16; Figure 5) and a A582V missense variant in *ADAM12* (QTLR on BTA26; Additional File 3: Figure S10). Note that these genes have multiple transcripts and the position of the amino acid change might thus vary (the genomic coordinates and alleles of the variants are available in Table 2). These three variants have strong statistical support (Additional File 1: Table S6). The two first missense variants were indeed the lead SNP in the MT GWAS whereas the third was almost in perfect LD (r² = 0.998) with the lead SNP of the MT GWAS for muscular development traits (Additional File 3: Figure S10). In addition, each of these variants was also the lead SNP and present in both SuSiE and LD-based CS for at least one ST GWAS.

**Figure 5.**
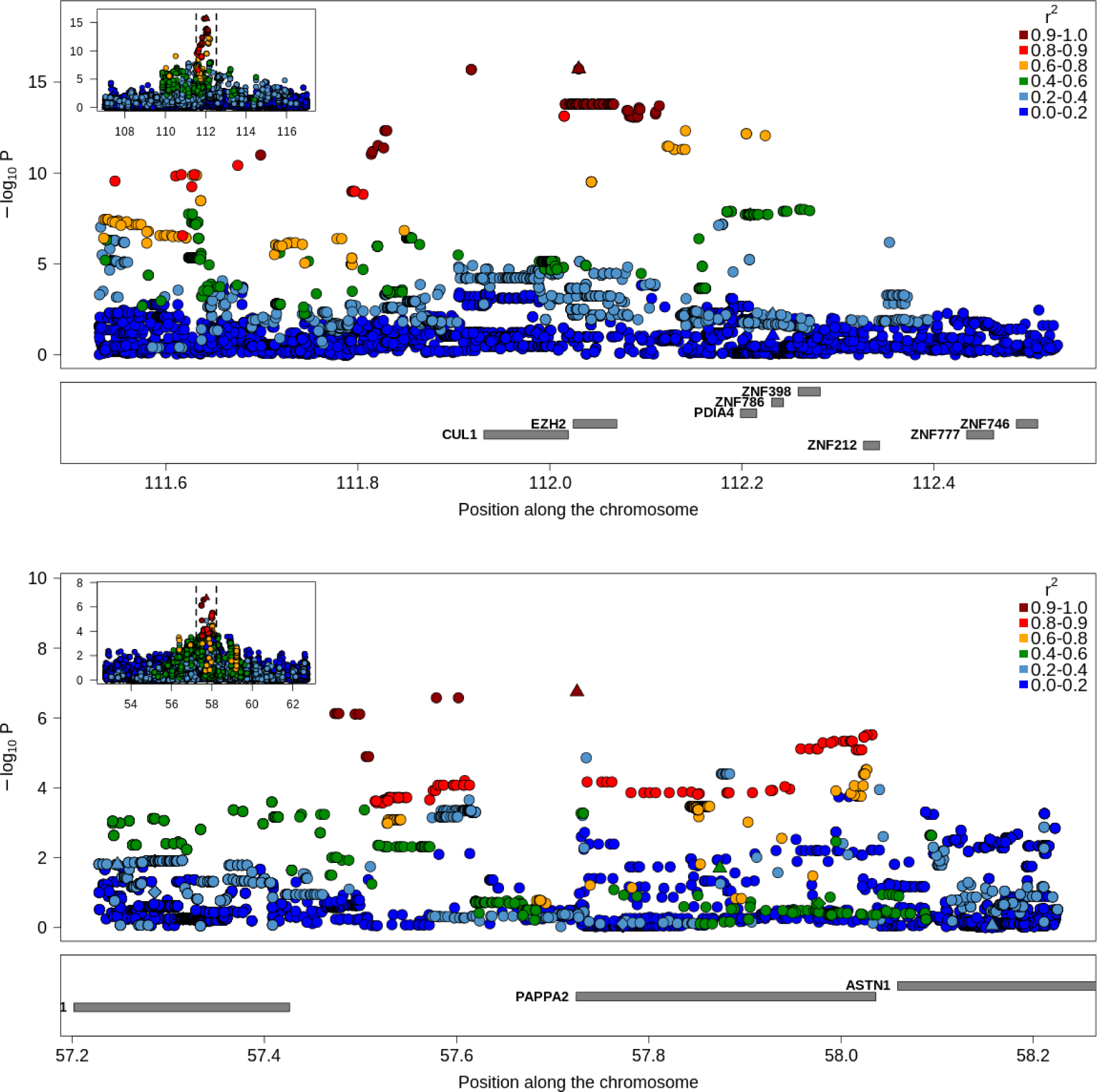
Regional association plot for the QTLR with missense variants in *EZH2* and *PAPPA2*. The colors represent the LD level with the lead variant and the symbols indicate the predicted impact of the variant (● modifier, ♦ low impact, ▴moderate impact, ▪ high impact). The positions of the genes are in the lower track. Upper panel: results of the MT-GWAS with traits related to body size on BTA4 and encompassing *EZH2*; lower panel: results of the MT-GWAS with traits related to muscular development on BTA16 and encompassing *PAPPA2*.

In total, six coding variants were identified in the 11 MT LD-CS associated with six QTLR (when using a LD threshold of 0.90). Using the proportion of variants with moderate or high predicted impact (mainly missense, splice site and frameshift variants and premature stop codons) in our data (0.34%) and the size of each CS, we estimated by random sampling (10^8^ repetitions) that we expect only 1.1 such variants on average in our CS. The enrichment of these variants in our CS was significant (p = 0.001) and the number of CS harboring at least one coding variants was also significantly higher than expected (p = 2.0e-6). The chance to have a coding variant as lead variant for five or more CS was even lower (p < 1e-8). If we define CS using a 0.80 LD threshold, three additional coding variants would be identified including a frameshift variant in *LCORL* (a 2bp deletion ACT > A at position 37401770) included in a long LD block with 73 variants spread over 2 Mb (see Additional File 3: Figure S11). This would lead to a total of 9 coding variants located in 8 distinct QTLR regions (MT LD-based CS containing then on average 73 variants). The number of observed coding variants and the number of CS harboring at least one coding variants would still be significant (p = 2.1e-3 and 3.0e-6).

### Evidence for regulatory variants among QTLR

For the QTLR without obvious candidate coding variant, we also investigated whether SNP in the CS overlapped with core and consensus segments from ATAC-seq peaks present in the catalogue from Yuan *et al*. (2023) or with cis-eQTL reported in the same study. Contrary to the lead variant on BTA6, the CS variants on BTA5 fall in consensus ATAC-seq peaks and even in the CS from a liver eQTL (the lead SNP matches both criteria, the alternate allele being associated with increased expression and higher height). Interestingly, a SNP located at position 105,773,809 (A>G) and in LD with the lead SNP (r² = 0.89) was the lead variant from a meta-analysis involving 18 breeds [8] and subsequently identified as a significant trans eQTL across multiple genes and tissues [50]. On BTA18, several CS variants were located in both consensus and core ATAC-seq peaks. Finally, in the CS for the QTLR on BTA23 containing only six SNP, four intergenic variants match core ATAC-seq peaks and are in the CS from a blood eQTL reducing *ARMC12* expression levels. The lead SNP located upstream from *ARMC12* is the lead SNP of this eQTL, the alternate allele is associated with lower expression and lower height.

### Several credible sets encompass genes associated with height in other species including *CCND2* and *LCORL*

Among QTLR with candidate coding variants, three were related to genes linked with height or growth in other species (*EZH2*, *PAPPA2*, *ADAM12*). Two of the remaining QTLR harbor strong candidate genes known to control height in other breeds or species. In the QTLR on BTA5 (Additional File 3: Figure S12), all the SNP in the CS were intronic variants from *CCND2*. On BTA6, the lead SNP from the MT GWAS was an intergenic variant but *LCORL-NCAPG* were located in the same region (Additional File 3: Figure S11). In addition to these two genes previously reported for their association with height, *ARMC12*, the gene included in the CS for the QTLR on BTA23 (Additional File 3: Figure S13), presents also some link with height. Indeed, this gene increases the activity of *EZH2* [51], our candidate gene for the QTLR on BTA4. Note that we did not find evidence for interactions among the two identified variants (*i.e.* the effect of the variant in *EZH2* is the same when estimated conditionally on the three possible genotypes at the *ARMC12* variant). Variants in *ARMC12* have been recently associated with human height in a large-scale GWAS [52]. Finally, the CS on BTA18 still contained 86 variants associated with 12 genes (Additional File 3: Figure S14) and no obvious candidate variant could be defined. However, associations were found in the human orthologous region in the same GWAS [52]. In total, six of our QTLR can be linked to genes affecting height in other species.

### Stepwise conditional mapping: independent associations in *CCND2* and *LCORL* and association with an additional deleterious coding variant

For each QTLR, we selected variants to fit as covariate in a secondary mapping analysis. In QTLR with candidate coding variants, we chose these as they were excellent functional candidates and presented very strong statistical significance (e.g., lead SNP in MT GWAS). For the other QTLR, we used the lead SNPs for subsequent analysis (Table 2). We performed the conditional mapping in 10 Mb regions centered around the selected variants. As for the initial mapping, CS are reported in Additional File 2. For the six QTLR on chromosomes 3, 4, 14, 16, 18 and 26, no new significant associations were detected with the conditional mapping (Figure 6 and Additional File 3: Figure S15 – Table S7), indicating that the fitted variant captured the QTL signal for all associated traits. For many of the QTL or QTLR, the signal dropped strongly (see for instance examples on BTA3, BTA4, BTA14 or BTA26). However, for the two QTLR regions mainly association with body dimension traits and located on BTA5 and BTA6, new significant associations with the same group of traits were detected (Figure 6 and Additional File 3: Figure S16). These QTLR presented among the most significant associations in the first scan, and still harbor highly significant associations (p < 1e-7 and 1e-8, respectively). On BTA5, the exact same group of three traits was associated (size, body length, pelvis length) and all the CS encompass a single SNP downstream of *CCND2*. On BTA6, the association was significant for size and body length. The MT LD-based CS was particularly large, including several variants in *NCAPG* or *LCORL* (e.g. intronic) embedded in a long haplotype block (see Figure 6). Among those, the variant with the largest predicted impact was a missense variant Y551C in *LCORL*, presenting a r² = 0.95 with the lead SNP. The LD between the lead SNP from the primary and secondary associations were low, respectively 0.16 and 0.01 for QTLR on BTA5 and BTA6, indicating independent associations. For both these QTLR, the conditional mapping identifies thus a second independent QTL associated with the same set of traits and pointing to the same genes.

**Figure 6.**
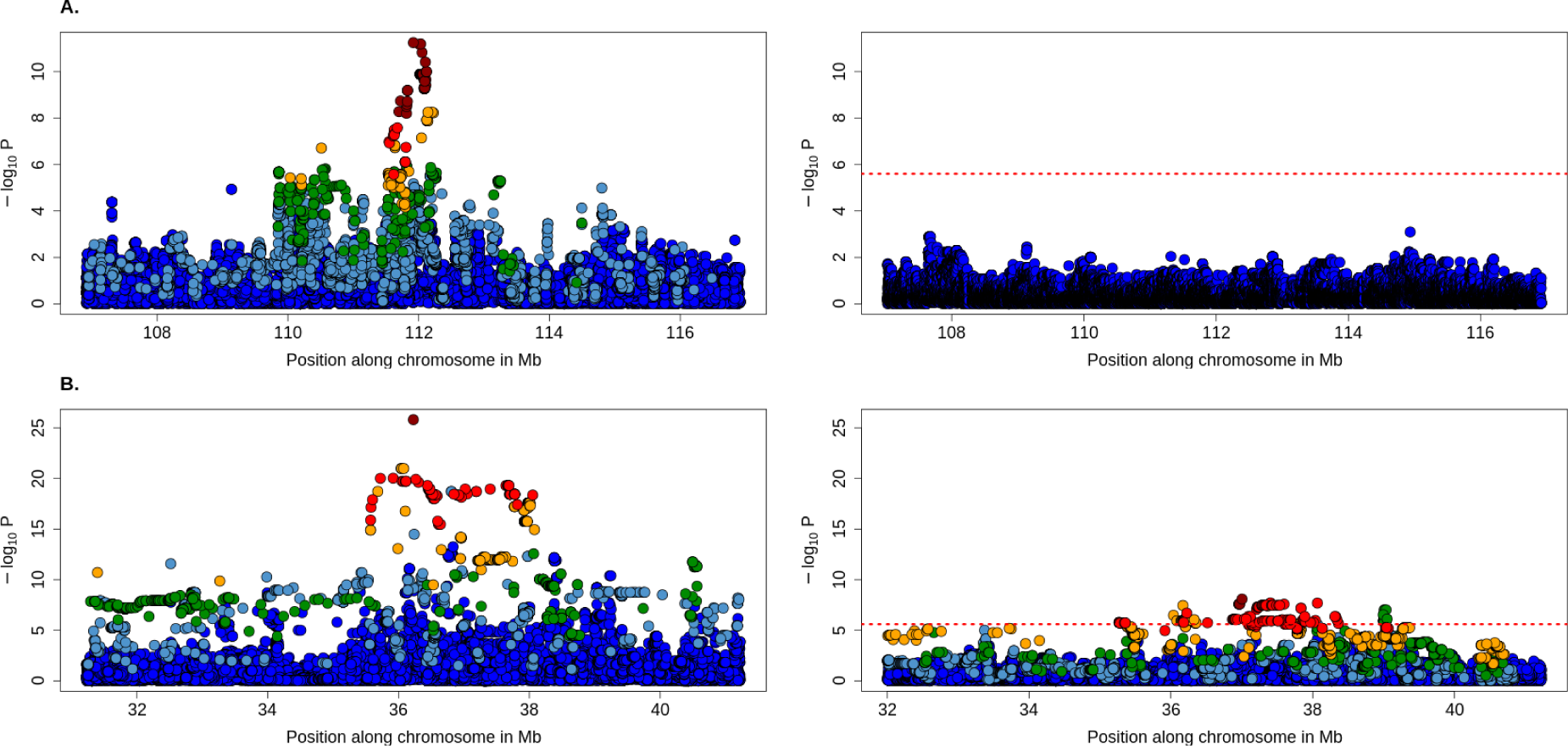
Regional association plot for the conditional mapping in QTLR on BTA4 and BTA6. The left panels represent the initial GWAS whereas the right panels correspond to conditional GWAS in which the candidate variants are fitted as covariate. The colors represent the LD level with the lead variant. The positions of the genes are in the lower track. **A**) GWAS for pelvis width on BTA4, and **B**) GWAS for length on BTA6.

For the three last QTLR regions, significant associations were still present but at lower magnitude (p > 1e-7). These signals would not be significant for a whole-genome scan but indicate that all the primary signal has not been captured by fitting our candidate variants. For the QTLR on BTA19, the most significant signals drop after inclusion of the LoF variant in *MRC2* (Additional File 13 Figure S17). However, associations are significant for body length and top muscling (-log_10_P > 5.6) whereas there is still some evidence for association with size or rump (p < 1e-5). For body length and top muscling, the CS contains a single SNP (respectively, an intronic variant in *MAPT* and an intergenic variant). These four associations point to four different regions, indicating a quite complex QTLR. On BTA23, the lead variant, associated with *ARMC12*, captured the signal for body length and pelvis length whereas for size, there was signal for a second QTL (p = 2.4e-6) located at more than 2 Mb (Additional File 3: Figure S16). The lead SNP was an intron variant in *BOLA-DOB* (the CS contained only one more SNP). To note, this is a complex region with high diversity and low LD levels. Finally, for the QTLR on BTA25, there was no longer evidence for association with rump after inclusion of the variant in *ATP2A1* in the model (Additional File 3: Figure S18). However, this variant did not capture the signal associated with muscular development of the buttock (side view) for which the association was still strong (p = 1.35e-7; Additional File 3: Figure S18). There is thus evidence for two linked QTL affecting two distinct traits in that QTLR. Two distinct and distant peaks achieved similar significance levels (Additional File 3: Figure S18; Additional File 2), the CS for the first peak consisted in a single intergenic variant whereas the second CS contained 15 SNP (r² > 0.90). A R631W missense variant in *ATP2A1* previously shown to affect negatively meat quality and muscular development [53] was in high LD (r² = 0.88) with the lead variant and represents thus the best candidate causative variant.

We repeated a conditional mapping by adding the lead variants for the secondary QTL on BTA5, BTA6, BTA23 and BTA25. For all tested traits and QTLR, we did not detect new significant associations after correction for multiple testing (see Additional File1: Table S8). The p-values were indeed higher than 1e-4, whereas the significance threshold was set to 2e-5 (considering ∼2500 independent SNP in the four tested QTLR).

## Discussion

We herein performed a sequenced-based association study for 11 traits related to muscular development and body dimension in a cohort of ∼15,000 BBB cattle cows. We identified 11 QTLR with genome-wide significant associations and most of them affected several correlated traits. Several coding variants included in our CS represented strong candidate causative variants. Five of these correspond to deleterious variants specific to BBB that have been previously characterized [18–20, 49, 53]. In addition, we found three new missense variants in *PAPPA2*, *ADAM12* and *EZH2*, three genes related to growth disorders in different species including human. Indeed, the role of *PAPPA2* on growth has been documented in multiple species, it is a regulator of *IGF1* and is associated with short stature in human [54, 55] and in mice [56, 57]. *ADAM12* was identified as a susceptibility gene for Kashin-Beck disease, causing growth retardation [58]. In agreement, *ADAM12*-deficient zebrafish present growth retardation [59]. In human, mutations in *EZH2* cause the Weaver syndrome and increased height [60], tall stature [61] but also growth retardation and severe short stature [62]. Two of these coding variants were the lead variants in their respective MT-GWAS. They make thus strong candidates as they have strong statistical support, they change the protein sequence, and coding variants in the same genes are known to affect growth in other species. The three variants are predicted to be deleterious (SIFT score < 0.05) and have high PhastCons scores (> 0.88) and positive, although not extreme, GERP scores (from 0.22 to 1.22). In addition, two independent signals on BTA6 might be associated to a missense and a frameshift variant in *LCORL*, a gene associated with height in different cattle breeds [8] and several species. To our knowledge, these are the first coding variants in *LCORL* significantly associated with height reported in cattle. The Y551C missense variant in *LCORL* was predicted to be tolerated (SIFT score = 0.46) and was not conserved (0.00 PhastCon; –2.37 GERP score). In both cases, the variants were included in a long haplotype block encompassing many variants making it more difficult to pinpoint the causative variant. In addition, the frameshift variant was not in very high LD with the lead SNP (0.84). The evidence for their causality is thus weaker, although they might affect the protein function of a strong candidate gene. Overall, the number of coding variants in our CS and the number of CS harboring at least one coding variant were significantly higher than expected by chance. These enrichments suggest that a fraction of these coding variants are causative, in particular if we consider that several of them were lead SNP (which is even less likely by chance) and that they fall in genes previously associated with height. The number of QTL is too limited to make strong assumptions on the relative contribution of coding versus regulatory variants to genetic variation of complex traits. Our QTL represent only a fraction of variants contributing to genetic variation, and correspond only to the largest effects segregating in the BBB cattle population (see Additional File 2). Nevertheless, contribution of coding variants should not be underestimated.

Beside these candidate coding variants, we found evidence for regulatory variants in three QTLR on BTA5, BTA18 and BTA23. CS from these three QTLR overlapped indeed with the catalogue of regulatory regions identified by ATAC-SEQ by Yuan et al. (2023). For QTLR on BTA5 and BTA23, there was also association between the CS with cis-eQTL from a study conducted in blood and liver in Holstein (Yuan et al., 2023). In addition, the BTA5 QTL CS contained a SNP previously proposed as candidate variant for a stature QTL and significantly affecting expression in trans in multiple tissues [50]. This illustrates how such catalogues can help to better understand mechanisms underlying identified QTL.

In 2011, Pryce et al. [6] concluded that genes associated with height in human control also stature in cattle. In agreement, Bouwman et al. [8] demonstrated that genes associated with height in cattle GWAS were enriched in genes also reported in human GWAS for the same trait. Biaty et al. [63] identified such shared genes by comparing associations found in human by Wood et al. [64] or by Yengo et al. [65] with those found in cattle by Bouwman et al. [8]. In our study, several candidate variants were also associated to genes previously associated with growth or height in cattle and in other species. First, the most significant QTLR located on BTA6, included *LCORL-NCPAG*, that have been linked with height in cattle based both on association studies [e.g., 8, 66] and signatures of selection [e.g., 8, 13]. Similar findings have been reported in other species including human, dog and horse [11, 12, 67]. In cattle, associations have been observed in several breeds. Second, the region on BTA5 was among the most significant regions and the CS included only one gene, *CCND2*. This gene has also been previously associated with height in other cattle breeds [8, 68–70], and in other species such as human [71, 72]. As in our study, the allele reported in human by Stenthorsdottir et al. [71] was regulatory (increasing both expression and height). For both QTLR on BTA5 and BTA6, we identified two independent signals stressing the importance of these genes and strengthening the causality of *CCND2* (it was twice the single gene present in the CS). Next, *ADAM12* and *PAPPA2* are both associated with growth disorders (see above) and have been identified as ‘shared’ genes by Biaty et al. [63]. *PAPPA*, a paralog of *PAPPA2,* has also been listed by Pryce et al. [6] as a gene affecting height in both human and cattle, and was proposed as candidate gene for size in horse by Petersen et al. [12]. Interestingly, the lead or candidate variants associated with *CCND2*, *LCORL*, *ADAM12* and *PAPPA2* are segregating in other breeds from the Run 3.0 of 1000 Bull Genomes Project [73] indicating that these variants are relatively old. Overall, these results confirm previous findings indicating that a set of shared genes contribute to genetic variation of height in mammals. We strengthened the evidence that these genes are causal in cattle based on statistical evidence (e.g., lead variants, limited number of genes in the CS, multiple independent associations for some genes) and by the identification of coding variants in *PAPPA2* and *ADAM12*. Such candidate coding variants with strong statistical support (e.g., present as lead SNP for at least one GWAS) were not previously reported among the shared associations. These genes appear thus associated with height in multiple breeds or species. As the catalogue of genes associated with height will increase in parallel with size of cohorts (see for instance Yengo et al. [52] that reported associations in 7,209 non-overlapping segments), the list of shared genes will automatically follow the same trend. Nevertheless, the identified shared genes remain relevant has the sharing was determined with relatively short lists of associated genes (i.e., before the study of Yengo et al. [52]). In addition, the four genes present multiple association in the human GWAS catalogues (https://www.ebi.ac.uk/gwas/): respectively 80, 26, 14 and 35 associations for *LCORL*, *CCND2*, *ADAM12* and *PAPPA2*. From these elements, we can thus conclude that these genes play an important role in genetic variation for height in multiple species.

For other QTLR, the candidate genes presented limited evidence for sharing across multiple species. For instance, *EZH2* is not associated with height in the cited GWAS catalogue, whereas *ARMC12* or the region on BTA18 encompassing genes such as *IL34*, *COG4*, *FUK*, *ST3GAL2*, *DDX19A* and *DDX19B*, present only associations in the largest cohort studies like those from Yengo et al. [52] or from Kichaev et al. [74]. In both studies, several genes are found in the associated genomic segments and the causative genes remains to be determined. As mentioned above, *EZH2* is nevertheless associated to growth disorders, and *ARMC12* is increasing *EZH2* activity [51]. These genes are involved in the polycomp repressive complex 2 (PRC2) that repress gene transcription during development through methylation [e.g., 75]. *EZH2* is the histone methyltransferase of PRC2 [e.g., 76, 77], whereas *ARMC12* facilitates the formation and activity of PRC2 [51]. These variants might thus play a role through epigenetic regulation. Unlike other identified genes, *EZH2* has not been reported in other GWAS for height. Interestingly, the missense variant is breed specific. Variants in this pathway seem thus to contribute less often to variation in height.

There was no obvious candidate gene in the CS on *BTA18* but we found evidence that regulatory variants might underlie this QTLR. Interestingly, the orthologous region in human is harboring enhancers. Such regulatory variants could influence other genes that overlap the QTLR. Among these *COG4* is a potential candidate gene as it is the causative gene for the Saul-Wilson syndrome causing dwarfism and skeletal abnormalities [78, 79], and is associated with reduced body length in zebrafish [80]. A mutant that affect skeleton and bone mineral density in mouse has been described in the Mouse Genome Informatics database [81]. Mutations in *COG4* have been show to disturb the Wnt signaling pathway playing an important regulation roles during embryonic development [80].

The four remaining QTLR were associated with five recessive deleterious variants previously identified in BBB cattle [18–20, 49, 53], including genetic defects [18, 19, 49]. Some of these variants present also heterozygous advantage [18, 19, 53]. These variants are breed specific (i.e., not observed in other breeds from the Run 3.0 of 1000 Bull Genomes Project [73]) and the associated genes, *RNF11*, *WWP1*, *MRC2* and *ATP2A1* are not associated to height in the GWAS catalogue (with the exception of *MRC2* that has associations in the large GWAS study from Yengo et al. [52]). These mutations have thus different properties and are quite specific to the BBB cattle breed that has experienced an intensive selection for muscular development, leading to the fixation of a 11bp deletion in the MSTN causing double muscling.

In conclusion, the BBB cattle breed represents an example of selection for extreme muscular development, mainly based on visual criteria. This led to the selection of variants associated with muscular development but also to selection for shorter stature. This process has resulted in the selection of several deleterious variants, some of which also associated to stature. Beside these deleterious variants, older variants present in several breeds and associated with genes affecting stature in multiple species also contribute to variation in height. The BBB cattle breed represents thus an interesting model to study height and identify new variants or implicated genes as *EZH2*.

## Supporting information

Credible Sets

## Acknowledgements

The authors acknowledge the Walloon Breeders Association (AWE group) for providing the data. Carole Charlier and Tom Druet are respectively Senior Research and Research Director from the F.R.S.-FNRS. We used the supercomputing facilities of the “Consortium d’Equipements en Calcul Intensif en Fédération Wallonie-Bruxelles” (CECI), funded by the F.R.S.-FNRS. The genotyping and sequencing were performed by the GIGA-Genomic platform.

