## Supplementary Tables for "Sequenced-based GWAS for linear classification traits in Belgian Blue beef cattle reveals new coding variants in genes regulating body size in mammals"

**Table S1.** Number of individuals genotyped on the different SNP genotyping arrays.

| Group of individuals | Number of individuals | Genotype array | | | | | Included in reference population | |
|---|---|---|---|---|---|---|---|---|
| | | Bovine LD | Illumina 50K | Eurogenomics MD | Bovine HD | WGS | MD reference | HD reference |
| Mapping population | 11,521 | x | | | | | | |
| Mapping population | 268 | x | x | | | | x | |
| Mapping population | 2,973 | | | x | | | x | |
| AI bulls | 66 | x | x | | | | x | |
| AI bulls | 658 | x | | | x | | x | x |
| AI bulls | 59 | | | | x | | | x |
| Individuals without phenotype | 133 | x | x | | | | x | |
| Individuals without phenotype | 9,502 | | | x | | | x | |
| Sequenced bulls (not in HD) | 173 | | | | | x | | x |
| Total | 25,353 | | | | | | 13,600 | 890 |

**Table S2.** Description of regions included in the conditional mapping analyses

| QTL-Region | Chromosome | Position on chromosome (in Mb) | Number of SNPs | Number of independent SNPs | Number of traits | Number of independent traits |
|---|---|---|---|---|---|---|
| 1 | 3 | 90.0 – 100.0 | 35,494 | 288.1 | 2 | 1.9 |
| 2 | 4 | 106.9 – 117.9 | 65,663 | 609.1 | 3 | 2.6 |
| 3 | 5 | 101.8 – 110.8 | 40,782 | 438.8 | 4 | 2.9 |
| 4 | 6 | 31.2 – 41.2 | 49,140 | 419.3 | 5 | 3.7 |
| 5 | 14 | 71.2 – 81.2 | 57,569 | 668.8 | 8 | 4.8 |
| 6 | 16 | 52.6 – 62.6 | 40,014 | 336.2 | 2 | 1.7 |
| 7 | 18 | 0.0 – 6.7 | 37,802 | 485.5 | 6 | 4.1 |
| 8 | 19 | 43.6 – 53.6 | 34,361 | 454.5 | 7 | 5.2 |
| 9 | 23 | 4.7 – 14.7 | 49,894 | 674.5 | 3 | 2.0 |
| 10 | 25 | 21.4 – 31.4 | 52,314 | 872.2 | 2 | 1.8 |
| 11 | 26 | 40.2 – 50.2 | 53,432 | 711.6 | 4 | 2.9 |

**Table S3.** Description of the 37 identified QTL. The frequency corresponds to the allele frequency of the ALT allele.

| QTL-Region | Trait | Position (bp) | REF | ALT | Freq | p-value |
|---|---|---|---|---|---|---|
| BTA3:95Mb | Top muscling | 95015373 | T | C | 0.05 | 4.35e-10 |
| BTA4:112Mb | Pelvis width | 111918080 | T | A | 0.47 | 5.69e-12 |
| | Height | 112030024 | T | C | 0.47 | 4.10e-10 |
| BTA5:106Mb | Length | 105762145 | C | G | 0.25 | 4.64e-16 |
| | Pelvis length | 105762145 | C | G | 0.25 | 2.69e-13 |
| | Height | 105762145 | C | G | 0.25 | 3.19e-13 |
| BTA6:36Mb | Pelvis width | 35683048 | G | A | 0.04 | 7.97e-10 |
| | Length | 36226849 | A | T | 0.05 | 1.55e-26 |
| | Pelvis length | 36226849 | A | T | 0.05 | 2.93e-16 |
| | Height | 36226849 | A | T | 0.05 | 2.41e-30 |
| | Top muscling | 37924938 | G | A | 0.05 | 3.28e-09 |
| BTA14:76Mb | Pelvis length | 75998203 | G | C | 0.16 | 7.05e-09 |
| | Length | 76227910 | C | T | 0.14 | 2.14e-10 |
| | Chest width | 76227910 | C | T | 0.14 | 5.13e-10 |
| | Pelvis width | 76227910 | C | T | 0.14 | 1.47e-11 |
| | Shoulder muscling | 76227910 | C | T | 0.14 | 4.63e-17 |
| | Top muscling | 76227910 | C | T | 0.14 | 6.30e-20 |
| | Buttock rear | 76227910 | C | T | 0.14 | 1.81e-19 |
| | Height | 76227910 | C | T | 0.14 | 6.85e-16 |
| BTA16:58Mb | Shoulder muscling | 57578954 | G | A | 0.72 | 3.35e-09 |
| | Chest width | 57725284 | C | A | 0.72 | 4.97e-09 |
| BTA18:2Mb | Buttock rear | 1165960 | C | CT | 0.26 | 2.03e-09 |
| | Chest width | 1673649 | A | AT | 0.27 | 3.59e-11 |
| | Shoulder muscling | 1717546 | G | A | 0.27 | 6.09e-09 |
| BTA19:49Mb | Top muscling | 47095175 | CAG | C | 0.03 | 1.15e-11 |
| | Rib shape | 47095175 | CAG | C | 0.03 | 8.85e-12 |
| | Buttock rear | 47095175 | CAG | C | 0.03 | 1.68e-09 |
| | Rump | 48297068 | T | C | 0.04 | 6.51e-09 |
| | Height | 48623853 | G | A | 0.03 | 7.21e-13 |
| | Length | 48707720 | C | T | 0.03 | 4.19e-09 |
| | Pelvis length | 48855548 | C | G | 0.04 | 1.70e-09 |
| BTA23:10Mb | Length | 9716619 | G | A | 0.48 | 9.30e-11 |
| | Pelvis length | 9716619 | G | A | 0.48 | 4.74e-09 |
| BTA25:26Mb | Rump | 26364983 | C | T | 0.01 | 1.68e-13 |
| BTA26:45Mb | Length | 45185751 | C | T | 0.62 | 2.47e-12 |
| | Pelvis length | 45489443 | T | C | 0.65 | 1.42e-09 |
| | Height | 45553105 | G | A | 0.59 | 9.13e-12 |

**Table S4.** Information on credible sets: number of traits sharing at least one variant in their credible set and number of trait with identical credible sets.

| QTL-Region | Number of traits | Number of traits with SNPs in common in the Credible Sets | | | Number of traits with identical SNPs in the Credible Sets | | |
|---|---|---|---|---|---|---|---|
| | | LD > 0.80 | LD > 0.90 | SuSiE | LD > 0.80 | LD > 0.90 | SuSiE |
| BTA3:95Mb | 1 | 1/1 | 1/1 | 1/1 | 1/1 | 1/1 | 1/1 |
| BTA4:112Mb | 2 | 2/2 | 2/2 | 2/2 | 2/2 | 2/2 | 0/2 |
| BTA5:106Mb | 3 | 3/3 | 3/3 | 3/3 | 3/3 | 3/3 | 3/3 |
| BTA6:36Mb | 5 | 5/5 | 3/5 | 5/5 | 0/5 | 0/5 | 0/5 |
| BTA14:76Mb | 8 | 8/8 | 7/8 | 8/8 | 0/8 | 0/8 | 0/8 |
| BTA16:58Mb | 2 | 2/2 | 2/2 | 2/2 | 0/2 | 2/2 | 0/2 |
| BTA18:2Mb | 3 | 3/3 | 2/3 | 3/3 | 0/3 | 0/3 | 0/3 |
| BTA19:49Mb | 7 | 7/7 | 5/7 | 7/7 | 0/7 | 0/7 | 7/7 |
| BTA23:10Mb | 2 | 2/2 | 2/2 | 2/2 | 2/2 | 2/2 | 2/2 |
| BTA25:26Mb | 1 | 1/1 | 1/1 | 1/1 | 1/1 | 1/1 | 1/1 |
| BTA26:45Mb | 3 | 2/3 | 0/3 | 2/3 | 0/3 | 0/3 | 0/3 |

**Table S5.** Comparison of LD-based credible sets (CS) and CS obtained with SuSiE. The threshold was set to 0.80 for the LD-based CS.

| QTL-Region | Trait | Number of SNP in LD-based CS | Number of SNP in SuSiE CS | Number of SNP shared between CS |
|---|---|---|---|---|
| BTA3:95Mb | Top muscling | 9 | 1 | 1 |
| BTA4:112Mb | Height | 72 | 30 | 30 |
| | Pelvis width | 72 | 30 | 30 |
| BTA5:106Mb | Height | 1 | 1 | 1 |
| | Length | 1 | 1 | 1 |
| | Pelvis length | 1 | 1 | 1 |
| BTA6:36Mb | Height | 74 | 1 | 1 |
| | Length | 74 | 1 | 1 |
| | Pelvis length | 74 | 1 | 1 |
| | Pelvis width | 74 | 30 | 28 |
| | Top muscling | 213 | 35 | 26 |
| BTA14:76Mb | Height | 276 | 101 | 101 |
| | Length | 276 | 121 | 121 |
| | Pelvis length | 174 | 174 | 132 |
| | Chest width | 276 | 132 | 131 |
| | Pelvis width | 276 | 67 | 67 |
| | Buttock rear | 276 | 34 | 34 |
| | Shoulder muscling | 276 | 81 | 81 |
| | Top muscling | 276 | 1 | 1 |
| BTA16:58Mb | Chest width | 105 | 13 | 13 |
| | Shoulder muscling | 120 | 2 | 2 |
| BTA18:2Mb | Chest width | 145 | 64 | 64 |
| | Buttock rear | 107 | 216 | 8 |
| | Shoulder muscling | 134 | 71 | 71 |
| BTA19:49Mb | Height | 22 | 1 | 1 |
| | Length | 24 | 1 | 1 |
| | Pelvis length | 10 | 1 | 0 |
| | Buttock rear | 29 | 1 | 1 |
| | Top muscling | 29 | 1 | 1 |
| | Rump | 16 | 1 | 1 |
| | Rib shape | 29 | 1 | 1 |
| BTA23:10Mb | Length | 13 | 6 | 6 |
| | Pelvis length | 13 | 6 | 6 |
| BTA25:26Mb | Rump | 24 | 8 | 8 |
| BTA26:45Mb | Height | 36 | 8 | 8 |
| | Length | 4 | 11 | 0 |
| | Pelvis length | 6 | 11 | 0 |

**Table S6.** Candidate or lead variants for the 11 QTLR. The table provides the number of times the variant was included in the CS (SuSiE and LD-based), the number of times it was the lead variant in ST-GWAS, and the LD between the candidate variant and the lead variant in the two MT-GWAS. MT-GWAS1: MT-GWAS with traits related to height and body dimensions; MT-GWAS2: MT-GWAS with traits related to muscular development.

| BTA | Position | Gene | Consequences | Presence in CS | | | Lead SNP in | LD with lead SNP | |
|---|---|---|---|---|---|---|---|---|---|
| | | | | SuSie | LD $r^2 > 0.90$ | LD $r^2 > 0.80$ | ST-GWAS | MT-GWAS1 | MT-GWAS2 |
| 3 | 95015373 | *RNF11* | Splice site variant | 1/1 | 1/1 | 1/1 | 1/1 | 1 | 1 |
| 4 | 112030024 | *EZH2* | Missense variant I549M | 2/2 | 2/2 | 2/2 | 1/2 | 1 | - |
| 5 | 105769735 | *CCND2* | Regulatory (ATAC-Seq, eQTL) | 0/3 | 0/3 | 0/3 | 0/3 | 1 | - |
| 6 | 36226849 | | Intergenic variant | 5/5 | 3/5 | 3/5 | 3/5 | 1 | 1 |
| 14 | 76227910 | *WWP1* | Missense variant R844Q | 7/8 | 7/8 | 7/8 | 7/8 | 1 | 1 |
| 16 | 57725284 | *PAPPA2* | Missense variant P282T | 1/2 | 2/2 | 2/2 | 1/2 | 1 | 1 |
| 18 | 1673649 | | Regulatory (ATAC-Seq) | 2/3 | 3/3 | 3/3 | 1/3 | 1 | 1 |
| 19 | 47095175 | *MRC2* | Frameshift variant | 7/7 | 3/7 | 6/7 | 3/7 | 1 | 1 |
| 23 | 9716619 | *ARMC12* | Regulatory (ATAC-Seq, eQTL) | 2/2 | 2/2 | 2/2 | 2/2 | 1 | 1 |
| 25 | 25933247 | *ATP2A1* | Missense variant R559C | 0/1 | 1/1 | 1/1 | 1/1 | - | 0.878 |
| 26 | 45553105 | *ADAM12* | Missense variant A582V | 1/3 | 1/3 | 1/3 | 1/3 | 0.711 | 0.998 |

**Table S7.** Most significant associations levels achieved for each trait in the conditional mapping. Association was performed only for traits presenting evidence for association in the first scan ($p < 1e-7$). Significance levels are expressed on a –log10 scale.

| QTL-Region | Length | Pelvis length | Pelvis width | Chest width | Shoulder muscling | Top muscling | Buttock side | Buttock rear | Rump | Rib shape | Height |
|---|---|---|---|---|---|---|---|---|---|---|---|
| BTA3:95Mb | 4.32 | | | | | 2.78 | | | | | |
| BTA4:112Mb | 3.19 | | 3.09 | | | | | | | | 3.46 |
| BTA5:106Mb | 7.04 | 5.88 | 5.37 | | | | | | | | 5.65 |
| BTA6:36Mb | 8.08 | 5.27 | 3.92 | | | 3.61 | | | | | 8.77 |
| BTA14:76Mb | 3.95 | 4.97 | 3.31 | 4.58 | 3.35 | 3.54 | | 3.60 | | | 5.02 |
| BTA16:58Mb | | | | 3.35 | 3.41 | | | | | | |
| BTA18:2Mb | | | | 3.73 | 3.38 | 4.45 | 3.25 | 4.00 | | | 3.08 |
| BTA19:49Mb | 5.91 | 4.31 | | | | 5.78 | | 4.70 | 5.46 | 3.72 | 5.07 |
| BTA23:10Mb | 3.92 | 3.46 | | | | | | | | | 5.62 |
| BTA25:26Mb | | | | | | | 6.82 | | 3.90 | | |

**Table S8.** Most significant associations levels achieved for traits in the second iteration of conditional mapping. Association was performed only for traits presenting evidence for association in the conditional mapping ($p < 2.0e-6$). The significance threshold was set at $–log_{10}P > 4.7$. Significance levels are expressed on a –log10 scale.

| QTL-Region | Length | Pelvis length | Buttock side | Height |
|---|---|---|---|---|
| BTA5:106Mb | 3.39 | 3.76 | | 3.57 |
| BTA6:36Mb | 4.03 | | | 2.77 |
| BTA23:10Mb | | | | 3.80 |
| BTA25:26Mb | | | 3.56 | |
