## Supplementary Figures for "Sequenced-based GWAS for linear classification traits in Belgian Blue beef cattle reveals new coding variants in genes regulating body size in mammals"

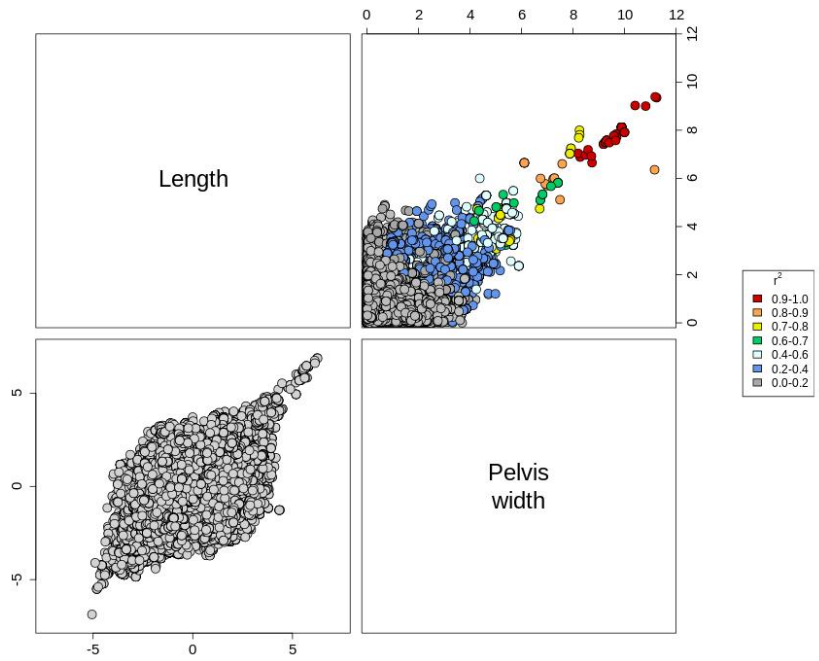


**Figure S1.** Scatterplots for association levels for different traits for the QTL region on BTA4. The selected traits are those harboring a significant signal in the QTLR. Upper diagonal: scatterplots with p-values on a negative log10 scale. The color represents the LD level with the lead SNP (from the trait with the strongest association). Lower diagonal: scatterplots with signed t-values.

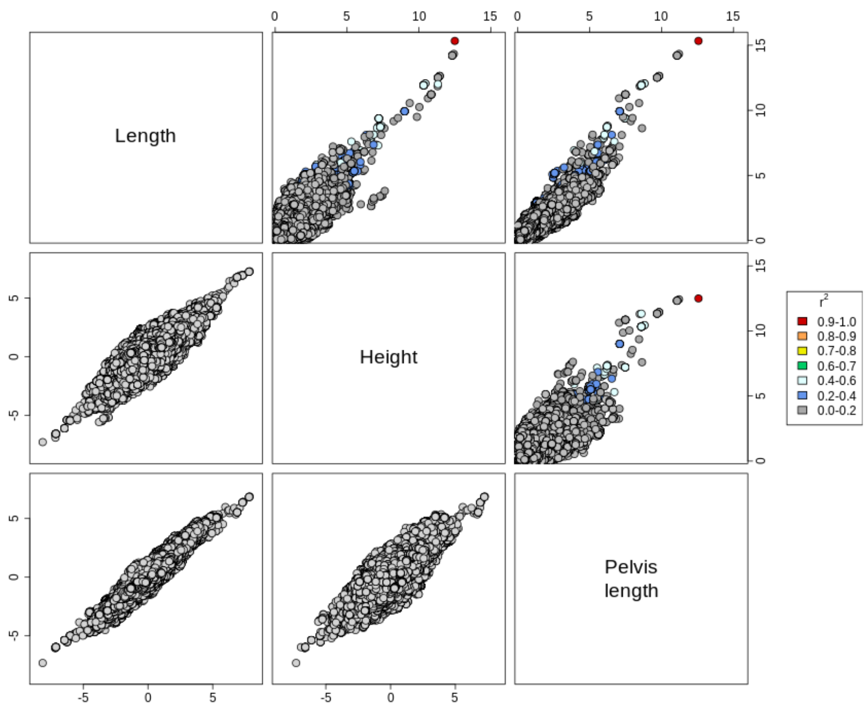


**Figure S2.** Scatterplots for association levels for different traits for the QTL region on BTA5. The selected traits are those harboring a significant signal in the QTLR. Upper diagonal: scatterplots with p-values on a negative log10 scale. The color represents the LD level with the lead SNP (from the trait with the strongest association). Lower diagonal: scatterplots with signed t-values.

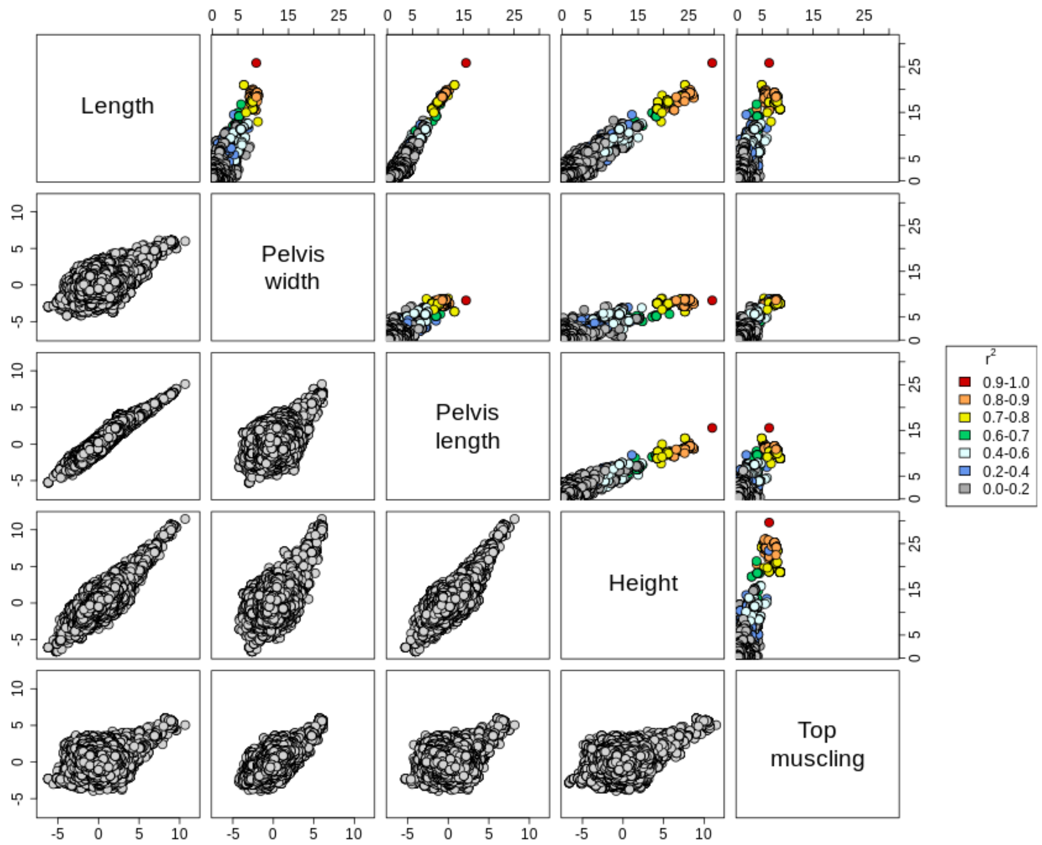


**Figure S3.** Scatterplots for association levels for different traits for the QTL region on BTA6. The selected traits are those harboring a significant signal in the QTLR. Upper diagonal: scatterplots with p-values on a negative log10 scale. The color represents the LD level with the lead SNP (from the trait with the strongest association). Lower diagonal: scatterplots with signed t-values.

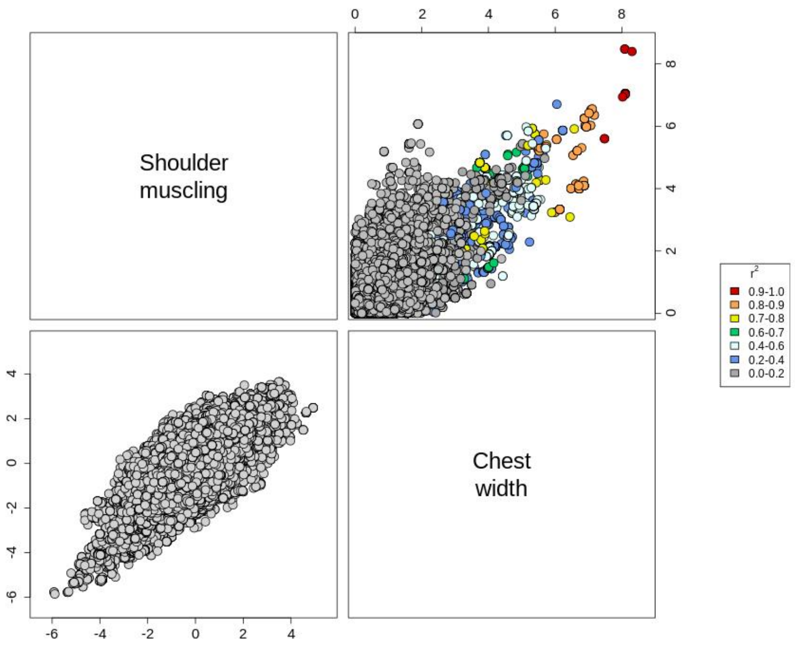


**Figure S4.** Scatterplots for association levels for different traits for the QTL region on BTA16. The selected traits are those harboring a significant signal in the QTLR. Upper diagonal: scatterplots with p-values on a negative log10 scale. The color represents the LD level with the lead SNP (from the trait with the strongest association). Lower diagonal: scatterplots with signed t-values.

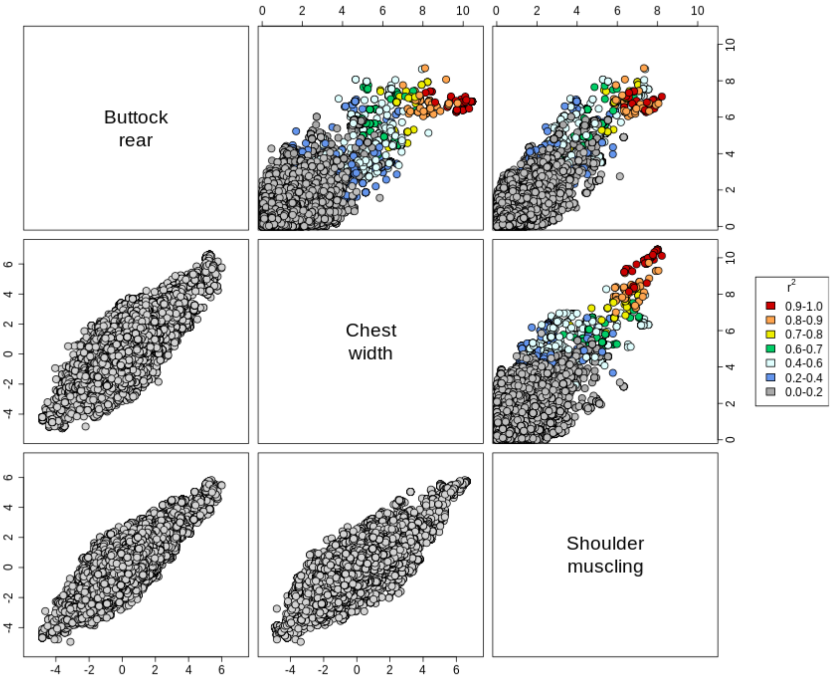


**Figure S5.** Scatterplots for association levels for different traits for the QTL region on BTA18. The selected traits are those harboring a significant signal in the QTLR. Upper diagonal: scatterplots with p-values on a negative log10 scale. The color represents the LD level with the lead SNP (from the trait with the strongest association). Lower diagonal: scatterplots with signed t-values.

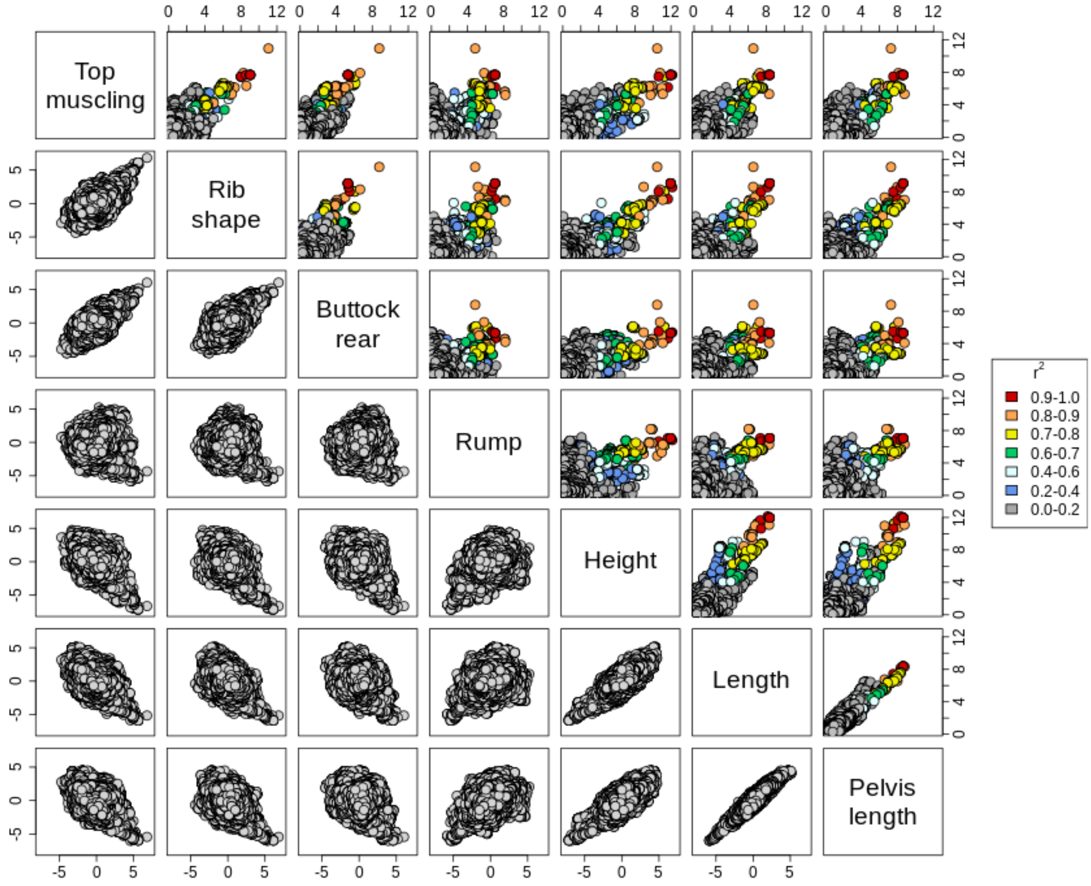


**Figure S6.** Scatterplots for association levels for different traits for the QTL region on BTA19. The selected traits are those harboring a significant signal in the QTLR. Upper diagonal: scatterplots with p-values on a negative log10 scale. The color represents the LD level with the lead SNP (from the trait with the strongest association). Lower diagonal: scatterplots with signed t-values.

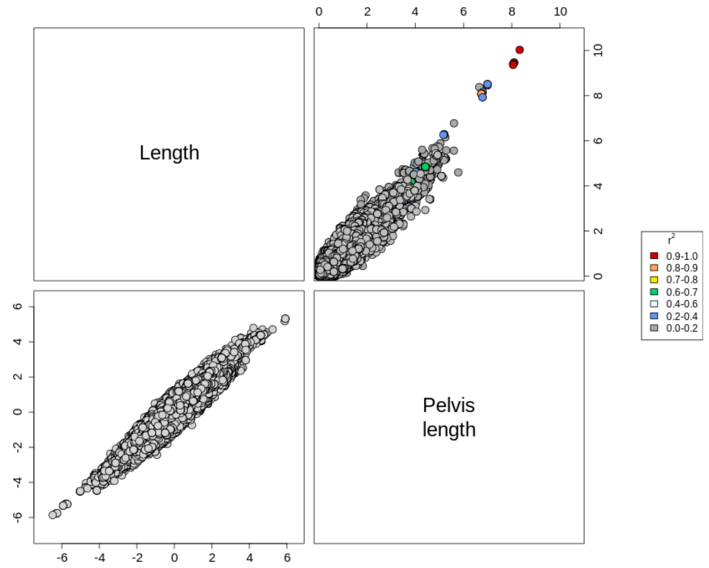


**Figure S7.** Scatterplots for association levels for different traits for the QTL region on BTA23. The selected traits are those harboring a significant signal in the QTLR. Upper diagonal: scatterplots with p-values on a negative log10 scale. The color represents the LD level with the lead SNP (from the trait with the strongest association). Lower diagonal: scatterplots with signed t-values.

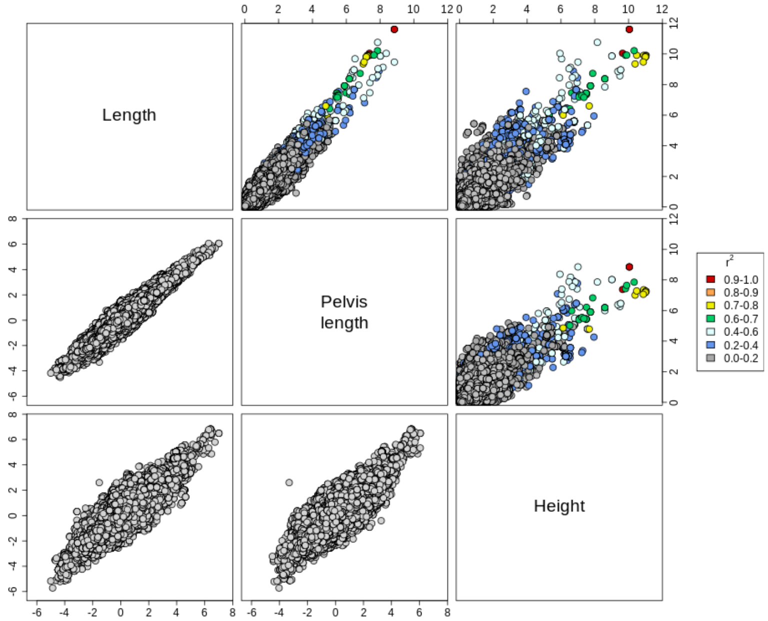


**Figure S8.** Scatterplots for association levels for different traits for the QTL region on BTA26. The selected traits are those harboring a significant signal in the QTLR. Upper diagonal: scatterplots with p-values on a negative log10 scale. The color represents the LD level with the lead SNP (from the trait with the strongest association). Lower diagonal: scatterplots with signed t-values.

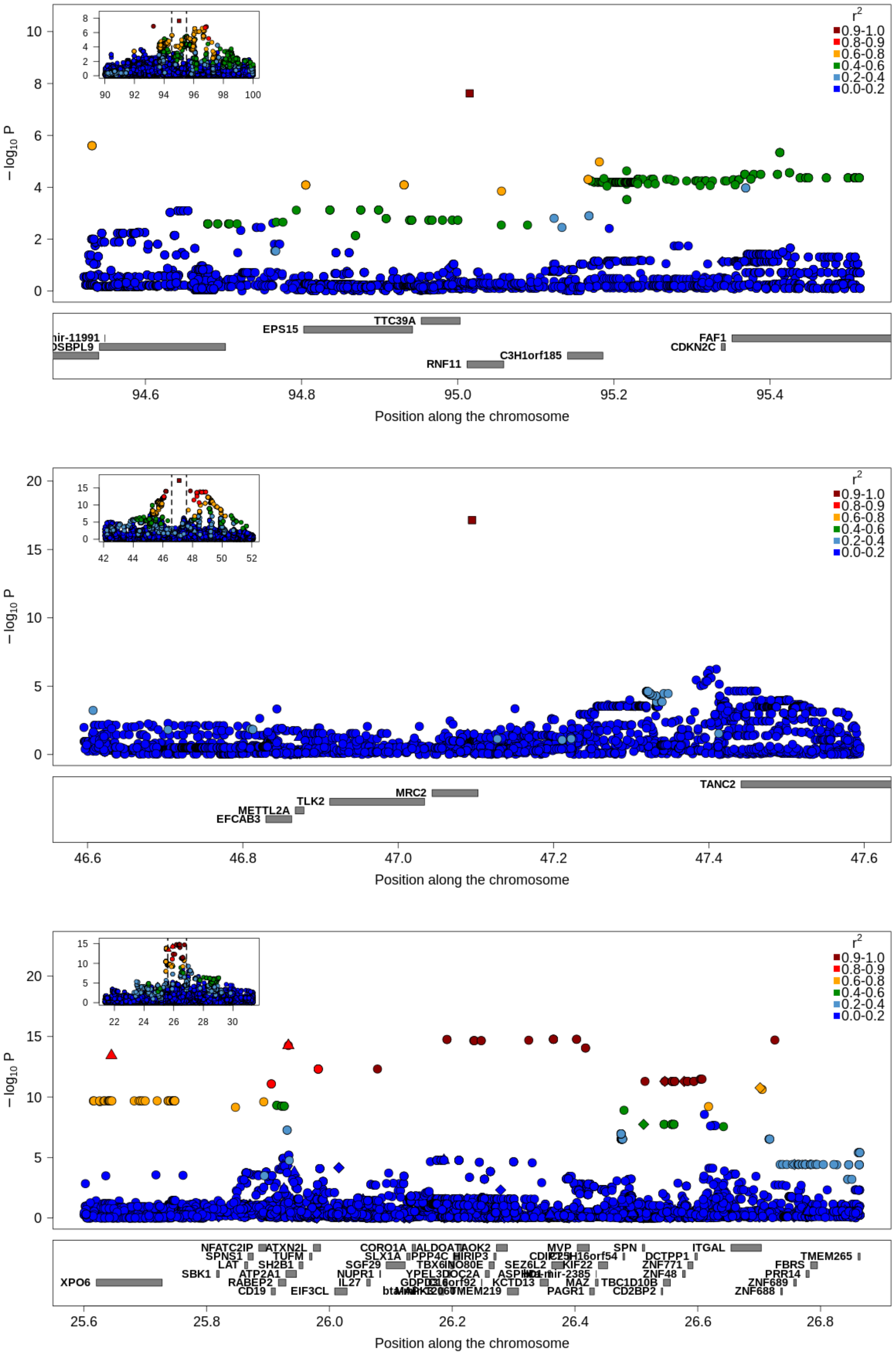


**Figure S9.** Regional association plot for the QTLR associated to recessive genetic defects. The colors represent the LD level with the lead variant and the symbols indicate the predicted impact of the variant (● modifier, ♦ low impact, ▲moderate impact, ■ high impact). The positions of the genes are in the lower track. Upper panel: mapping on BTA3 (MT-GWAS on muscular development traits); middle panel: mapping on BTA19 (MT-GWAS on body size traits); lower panel: mapping on BTA25 (MT-GWAS on muscular development traits).

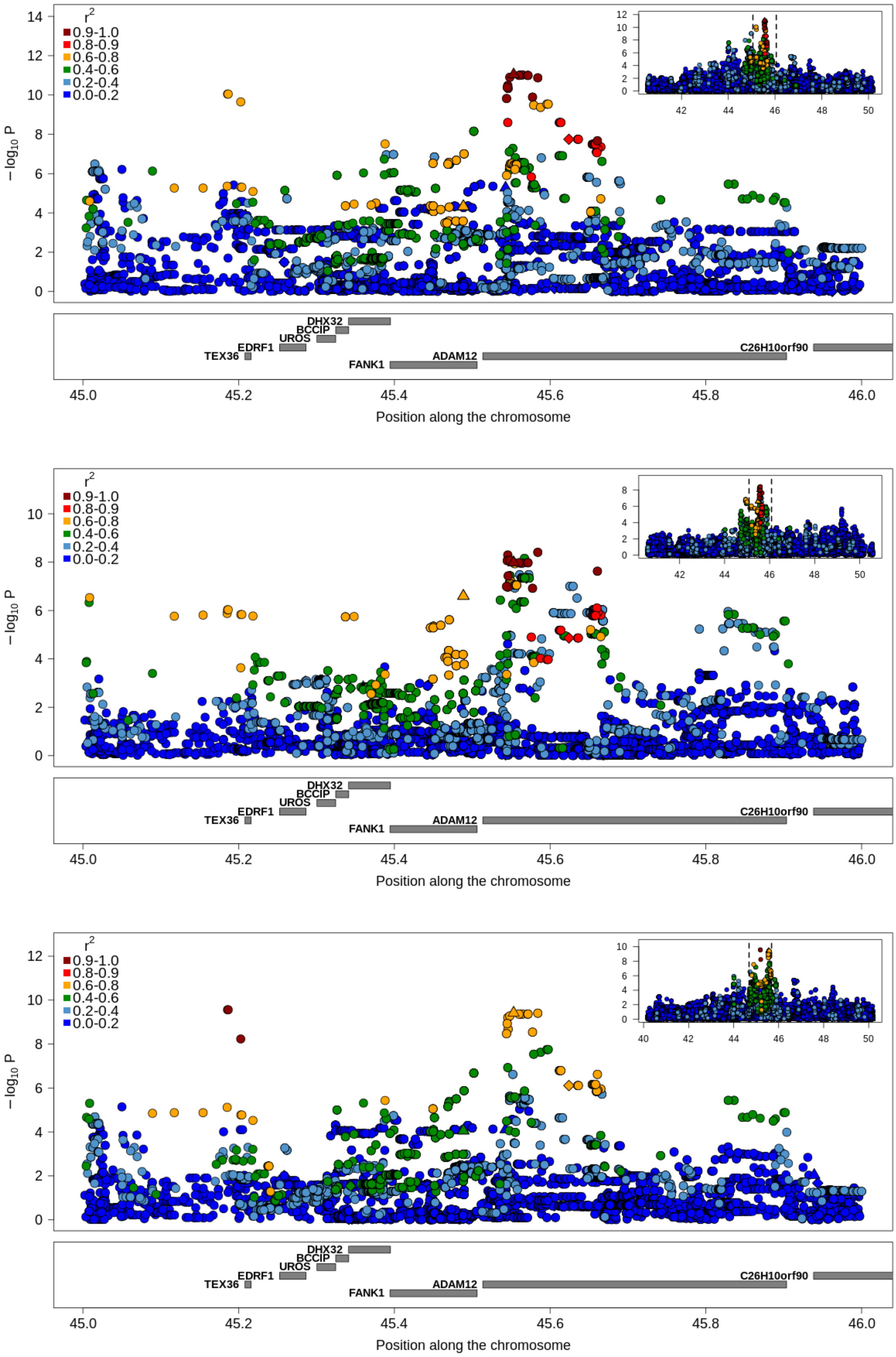


**Figure S10.** Regional association plot for the QTLR on BTA26. The colors represent the LD level with the lead variant and the symbols indicate the predicted impact of the variant (● modifier, ♦ low impact, ▲moderate impact, ■ high impact). The positions of the genes are in the lower track. Upper panel: ST-GWAS for height; middle panel: MT-GWAS on muscular development traits; lower panel: MT-GWAS on body size.

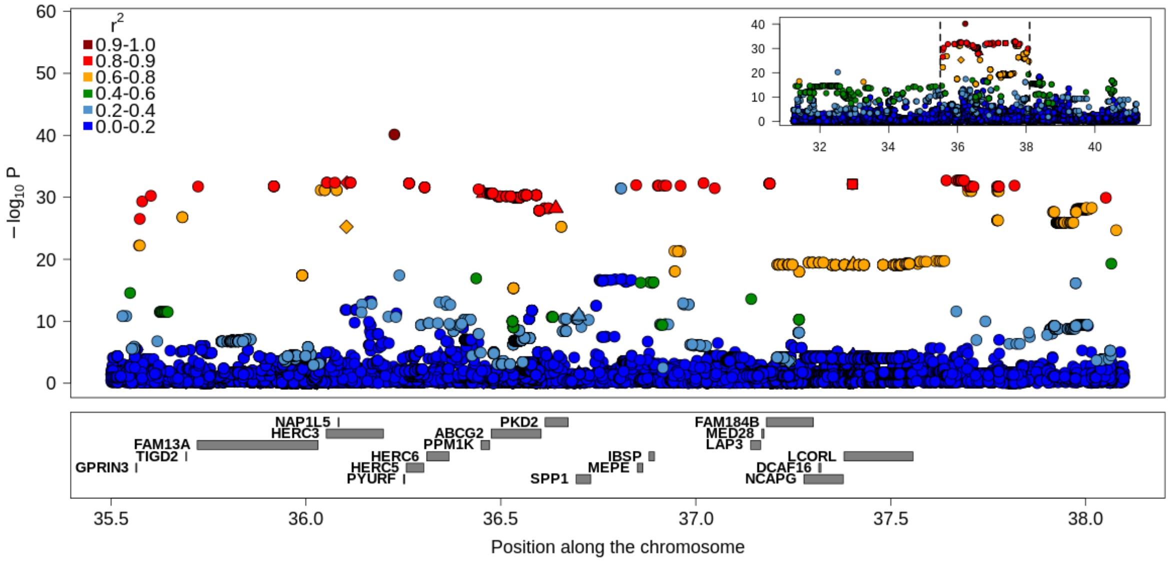


**Figure S11.** Regional association plot for the QTLR on BTA6. The results correspond to the MT-GWAS with traits related to body size. The colors represent the LD level with the lead variant and the symbols indicate the predicted impact of the variant (● modifier, ♦ low impact, ▲moderate impact, ■ high impact). The positions of the genes are in the lower track.

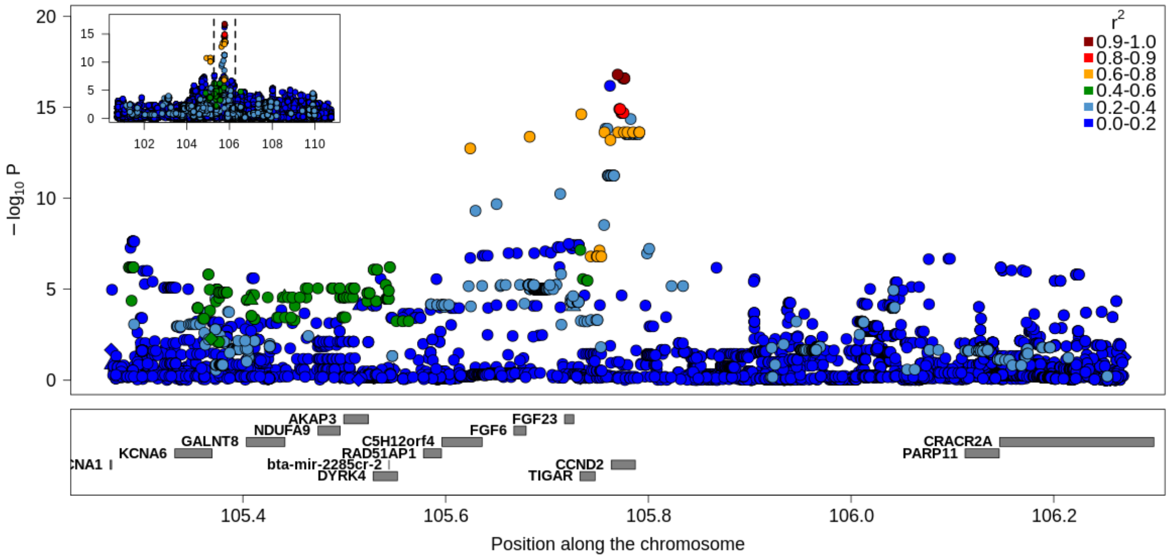


**Figure S12.** Regional association plot for the QTLR on BTA5. The results correspond to the MT-GWAS with traits related to body size. The colors represent the LD level with the lead variant and the symbols indicate the predicted impact of the variant (● modifier, ♦ low impact, ▲moderate impact, ■ high impact). The positions of the genes are in the lower track.

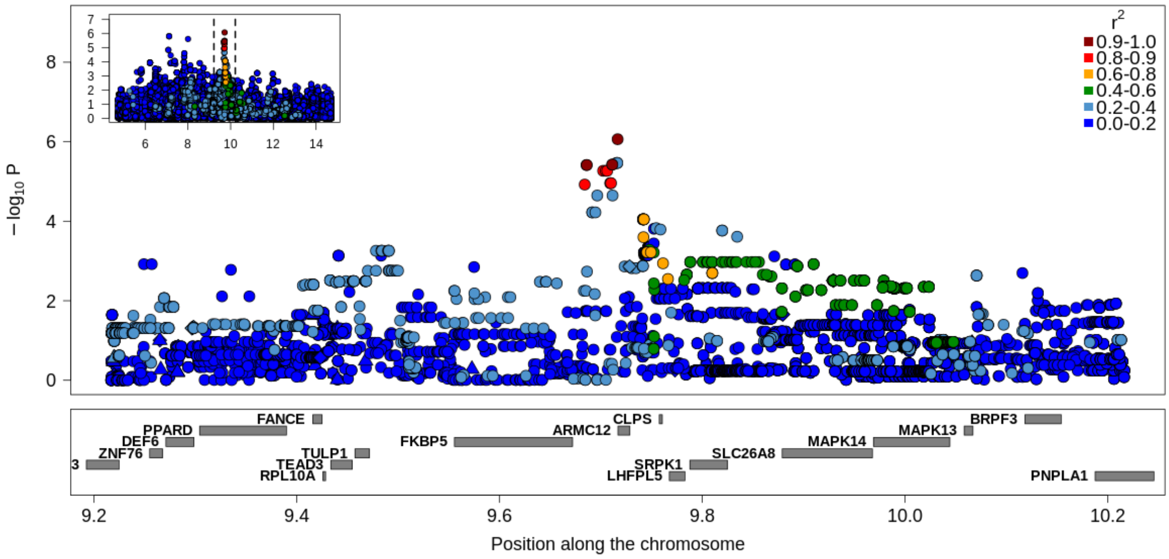


**Figure S13.** Regional association plot for the QTLR on BTA23. The results correspond to the MT-GWAS with traits related to body size. The colors represent the LD level with the lead variant and the symbols indicate the predicted impact of the variant (● modifier, ♦ low impact, ▲moderate impact, ■ high impact). The positions of the genes are in the lower track.

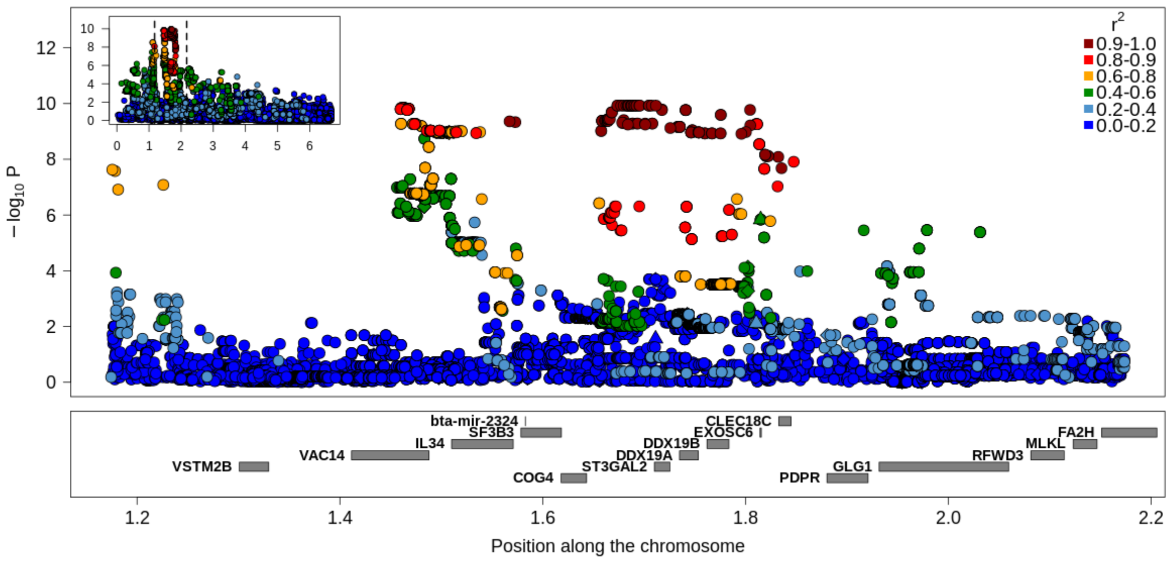


**Figure S14.** Regional association plot for the QTLR on BTA18. The results correspond to the MT-GWAS with traits related to body size. The colors represent the LD level with the lead variant and the symbols indicate the predicted impact of the variant (● modifier, ♦ low impact, ▲moderate impact, ■ high impact). The positions of the genes are in the lower track.

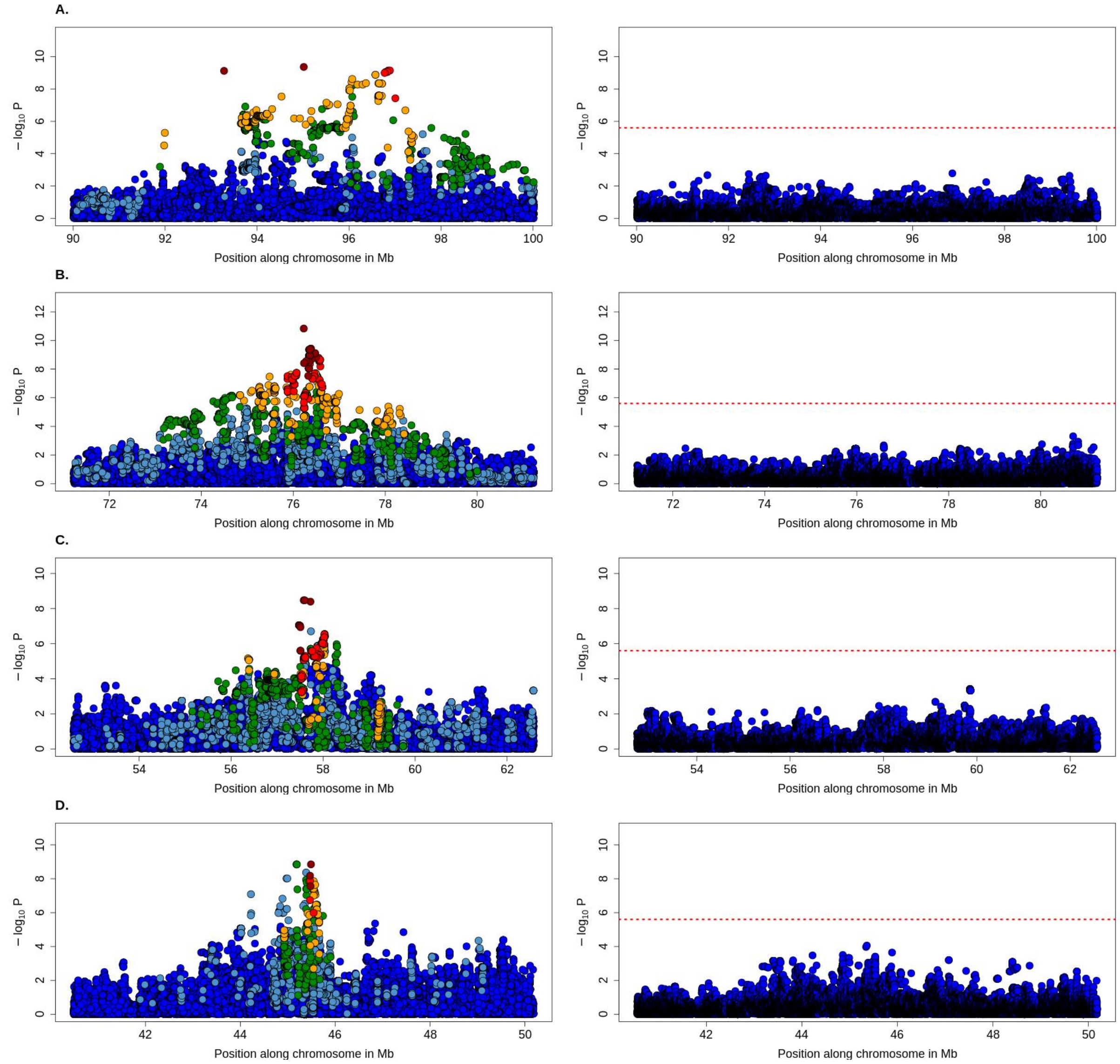


**Figure S15.** Regional association plot for the conditional mapping in QTLR on BTA3, BTA14, BTA16 and BTA26. The left panels represent the initial GWAS whereas the right panels correspond to conditional GWAS in which the candidate variants are fitted as covariate. The colors represent the LD level with the lead variant. The positions of the genes are in the lower track. **A**) GWAS for top muscling on BTA3, **B**) GWAS for pelvis width on BTA14, **C**) GWAS for shoulder muscling on BTA16, and **D**) GWAS for pelvis length on BTA26.

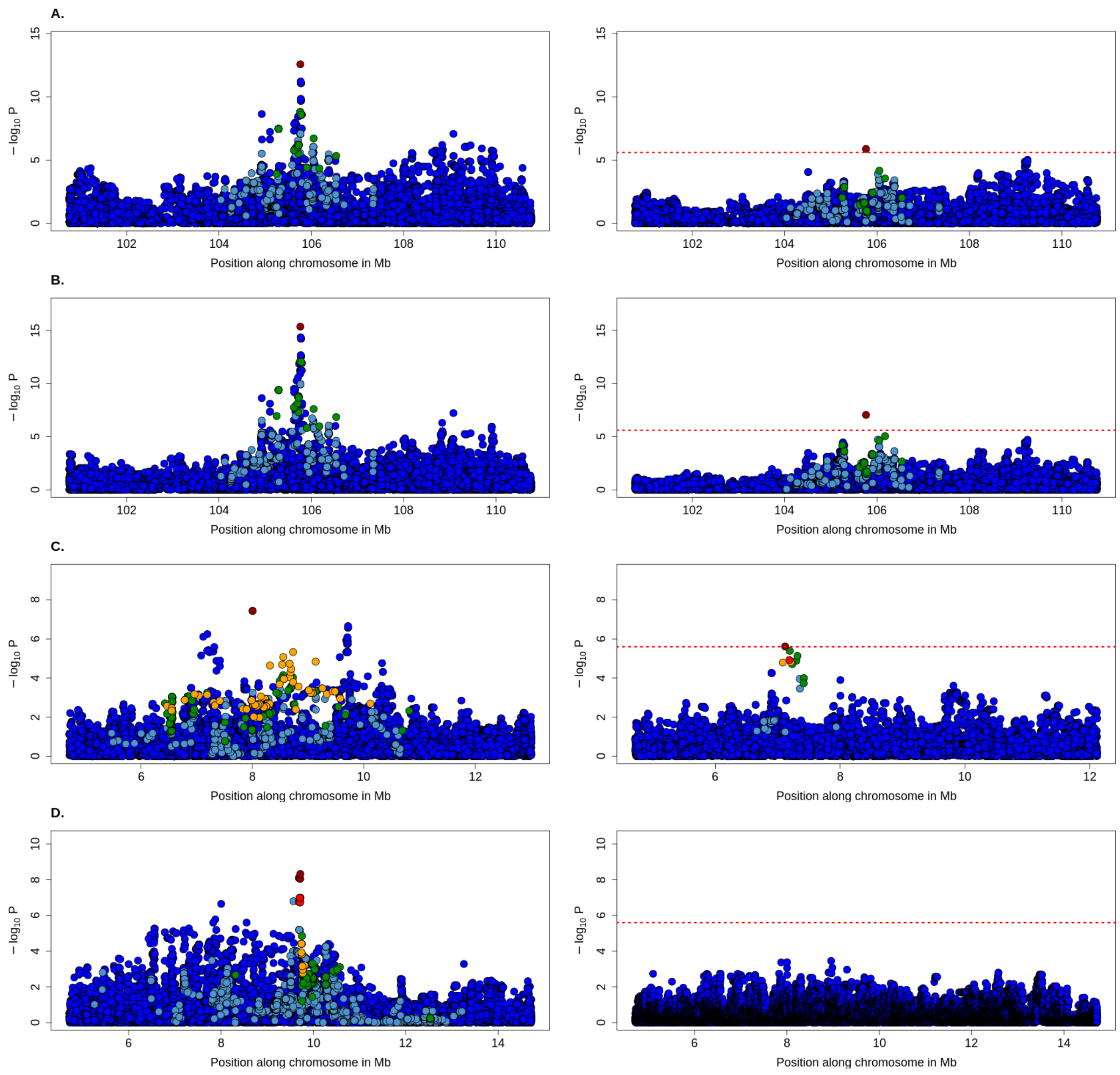


**Figure S16.** Regional association plot for the conditional mapping in QTLR on BTA5 and BTA23. The left panels represent the initial GWAS whereas the right panels correspond to conditional GWAS in which the candidate variants are fitted as covariate. The colors represent the LD level with the lead variant. The positions of the genes are in the lower track. **A**) GWAS for pelvis length on BTA5, **B**) GWAS for length on BTA5, **C**) GWAS for height on BTA23, and **D**) GWAS for pelvis length on BTA23.

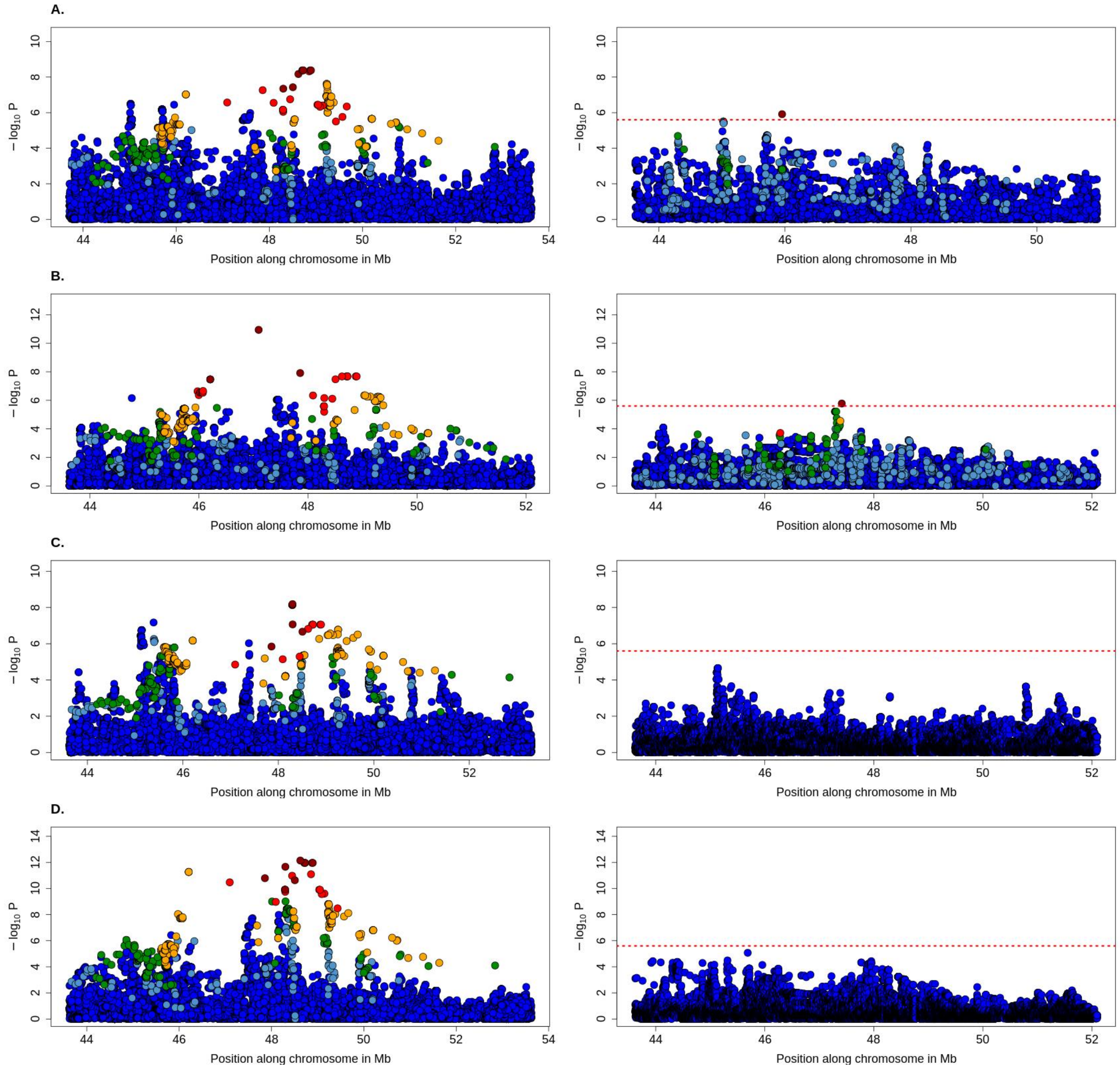


**Figure S17.** Regional association plot for the conditional mapping in QTLR on BTA19. The left panels represent the initial GWAS whereas the right panels correspond to conditional GWAS in which the candidate variants are fitted as covariate. The colors represent the LD level with the lead variant. The positions of the genes are in the lower track. GWAS for **A**) length, **B**) top muscling, **C**) rump, and **D**) height.

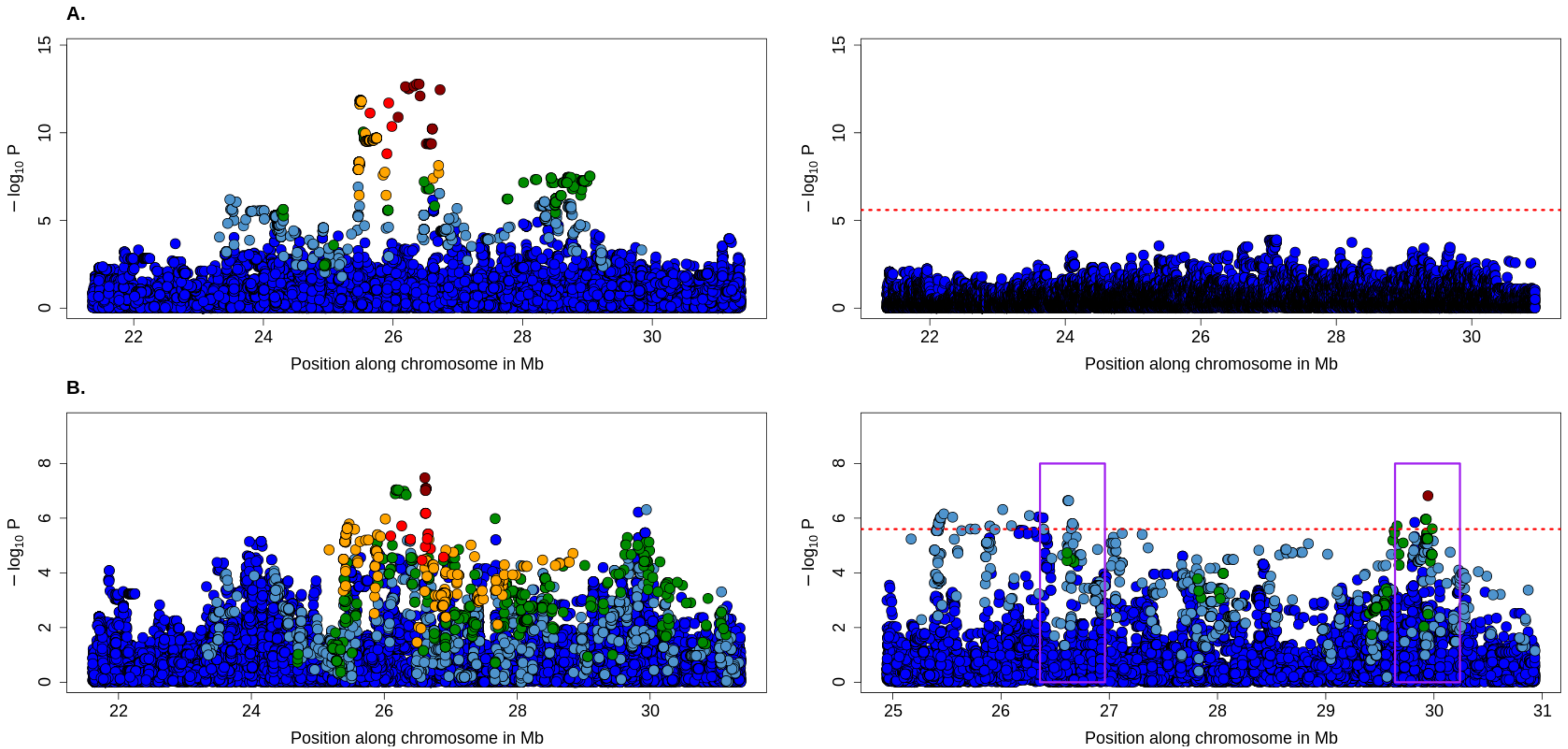


**Figure S18.** Regional association plot for the conditional mapping in QTLR on BTA25. The left panels represent the initial GWAS whereas the right panels correspond to conditional GWAS in which the candidate variants are fitted as covariate. The colors represent the LD level with the lead variant. The positions of the genes are in the lower track. GWAS for **A**) rump, and **B**) buttock muscling (side view). The boxes indicate the two regions achieving similar association levels and included in the credible set.

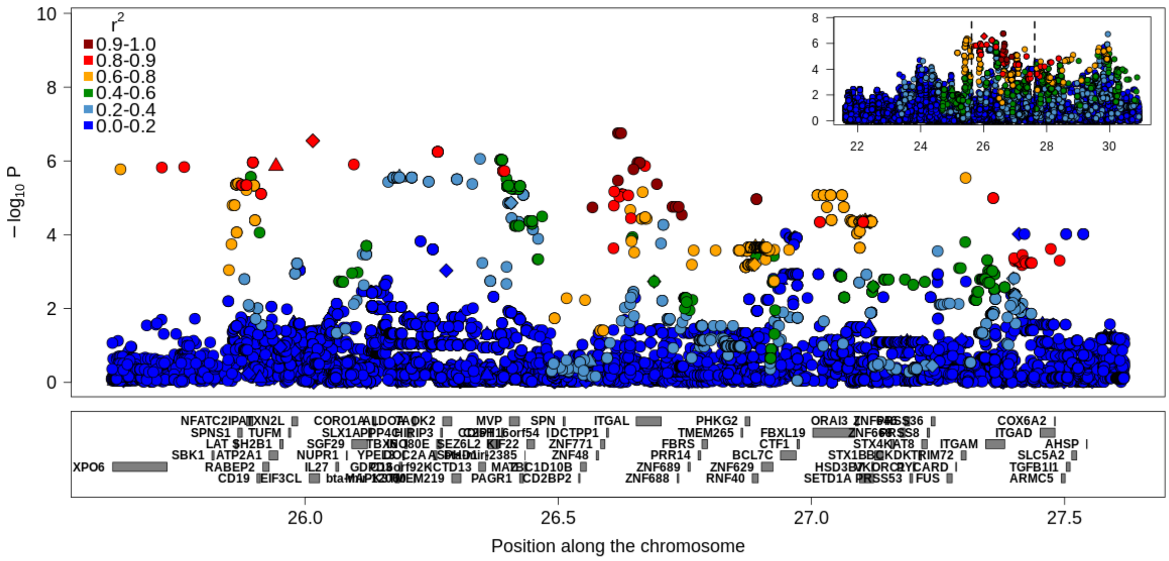


**Figure S19.** Regional association plot for the conditional mapping for the QTLR on BTA25. The results correspond to the MT-GWAS with traits related to buttock muscling (side view). The colors represent the LD level with the lead variant and the symbols indicate the predicted impact of the variant (● modifier, ♦ low impact, ▲moderate impact, ■ high impact). The positions of the genes are in the lower track.
